# Simple threshold-based Boolean rules fall short in capturing biological regulatory network dynamics

**DOI:** 10.1101/2025.06.26.661727

**Authors:** Priyotosh Sil, Olivier C. Martin, Areejit Samal

## Abstract

Among the various frameworks for modeling gene regulatory network (GRN) dynamics, Boolean modeling remains both powerful and accessible. Nevertheless, selecting update rules so that they capture the combinatorial control of various targets remains challenging due to limited quantitative data. Threshold majority rules (TMRs), a subtype of threshold functions (ThFs), update a gene’s state based on the signed sum of its regulators and its own activity, offering an elegant simplification. However, does the use of TMRs come at the cost of biological realism? Here, we rigorously evaluate the two standard TMR variants regarding their suitability in GRN modeling. We find that they provide limited canalyzation, exhibit discordant bias patterns, and often eliminate self-inhibitions. They are also underrepresented in empirical datasets, the opposite of what is expected for biologically relevant rules. Compared to nested canalyzing functions (NCFs), another class of ThFs but known to be preponderant in empirical datasets, TMRs exhibit heightened complexity and sensitivity to perturbations. At the network level, TMRs frequently fail to recover biological attractors and the associated basin size distributions. Using ensembles of random Boolean networks, we also show that one TMR type drives the network dynamics toward the chaotic regime. These findings invite a thoughtful reevaluation of TMRs as an appropriate logic for modeling GRNs despite their theoretical appeal.

## 1. INTRODUCTION

Genes coordinate essential cellular processes and guide the formation of diverse cell types during an organism’s development. To reveal the underlying actions, it is essential to understand how genes influence one another over time. Gene regulatory networks (GRNs) provide a unifying framework for describing cellular processes ranging from cell-fate decisions and apoptosis to proliferation and cell-cycle control [1–5]. Among the several modeling approaches for studying the dynamics of GRNs, the logical framework, particularly Boolean network (BN) modeling, has proven exceptionally useful. Its roots trace back to the work of Sugita [6, 7], who, inspired by Jacob and Monod [8], first applied logical circuits to gene regulation. Later, Kauffman [9, 10] and Thomas [11, 12] formalized this idea, demonstrating how a discrete ‘on/off’ representation of gene expression could capture essential features of network dynamics without requiring detailed kinetic parameters. In this approach, each gene or protein is assigned a binary state (‘on’ for active, ‘off’ for inactive) and a Boolean function (BF) which determines its next state based on the current state of its regulators. Kauffman initially used this framework to explore ensembles of random Boolean networks (RBNs), where both network topology and BFs were drawn at random [9]. Subsequently, in the genomic era, large-scale data-driven analyses have demonstrated that the architecture of biological networks exhibit significant deviation from randomness [13–15]. However, determining the precise regulatory logic (encapsulating the combinatorial control) at each node of a complex regulatory network remains a formidable task. As a result, modelers often rely on certain assumptions that can provide reasonable outcomes tailored to the specific context of their research.

The use of threshold functions (ThFs) in the context of modeling Boolean GRNs has drawn significant attention. The concept dates to the seminal work of McCulloch and Pitts, who modeled neuronal dynamics [16], and has since been widely adopted in the design of digital circuits and studies of artificial neural networks [17–20]. Surprisingly, the use of ThFs in the context of GRN modeling was initiated only in 1994 by Wagner, who focused on transcriptional networks [21, 22]. More recent work has shown that ThFs can be successfully applied to model the dynamical behavior of many different regulatory systems [23–26]. However, a realistic application of ThFs hinges on two types of parameters: the ‘weights’ or ‘strengths’ associated with the different regulators and the activation thresholds specific to each target node. Obtaining reliable values for these parameters is a major obstacle. For example, in their model on *A. thaliana* flower morphogenesis, Mendoza and Alvarez-Buylla treated both thresholds and interaction strengths as integers to simplify computations in the absence of detailed quantitative information [23]. Following a different strategy, Sevim and Rikvold [25] drew the interaction weights from a Gaussian distribution [25], while Li *et al*. [26] recently employed a regularized logistic regression to infer both the weights and the thresholds.

To circumvent the difficulty of parameter estimation, many modelers adopt a coefficient-free scheme where all regulatory interactions are assigned a strength of *±* 1 and activation thresholds are set to zero. Under these assumptions, the target node’s next state is solely determined by its own state and the weighted sum of its regulators’ current states. Two main variants of this approach have become standard in the literature. In the first variant, each node takes the values +1 (to indicate ‘on’) or *−* 1 (to indicate ‘off’). Throughout this manuscript we refer to this variant as IMR (Ising majority rule). This variant has been employed in the studies of epithelial-mesenchymal transition dynamics [27–29]. It has also been used in comparative analyses of frustration in biological versus random networks [30]. In the second variant, nodes take on binary values in the Boolean sense, adopting 1 for ‘on’ and 0 for ‘off’. We refer to this variant as BMR (Bit-based majority rule). This variant has been employed in the cell-cycle models of *S. cerevisiae* [31] and S. pombe [32]. BMRs have also found applications outside the realm of GRNs, for instance in modeling of the interactions of gut microbiome species [33, 34].

These simple variants (IMRs and BMRs) of ThFs, which we collectively refer to as threshold majority rules (TMRs), provide an elegantly simple and uniform framework for modeling GRNs. Nonetheless, it is important to examine if this minimalism overlooks key aspects of biological behavior. In this contribution, we probe the usefulness of TMRs based on features of BFs and models arising in empirical datasets and evaluate their viability as a go-to modeling strategy for GRNs. We begin by characterizing their basic properties like bias, effectiveness and canalyztion [35–37]. We then survey multiple empirical datasets of regulatory Boolean logic to explore how frequently these rules appear. Further, we compute their Boolean complexity [38] and average sensitivity [39], and compare these metrics to those of nested canalyzing functions (NCFs) [40], that are enriched in empirical datasets [37], to gauge TMRs’ intrinsic complexity and robustness to noise. Moving to the network level, we replace the BFs of the reconstructed models with the TMR variants and ask if this substitution still recovers the known biological attractors and their characteristic basin size distributions. Finally, we carry out a stability analysis across multiple measures and ensembles of RBNs to determine whether each variant steers the dynamics toward order or chaos. Altogether, through this comprehensive set of analyses, we show that although TMRs are very appealing from a theoretical point of view, they fail both at the single gene level (e.g., for combinatorial control) and at the network level (e.g., for reproducing the desired attractors).

## 2. METHODS

### 2.1. Boolean network modeling framework

A Boolean model of a biological regulatory network contains nodes (e.g., genes, proteins) that can assume one of two discrete states: active (‘on’, quantified as ‘1’) and inactive (‘off’, quantified as either ‘0’ or ‘-1’ depending on the framework) [10]. The nodes are connected by directed edges when there is a regulatory interaction between them which can be ‘activatory’ or ‘inhibitory’ or ‘mixed’. For a Boolean network (BN) with *N* nodes, we denote the state of the node *x*_*i*_ at time *t* by *x*_*i*_(*t*) for all *i ∈*{1, 2, …, *N*} and the state of the BN by **X**(*t*) = (*x*_1_(*t*), *x*_2_(*t*), …, *x*_*N*_ (*t*)). The temporal progression of the network is governed by a set of Boolean functions (BFs) {*f*_*i*_} or regulatory logic rules along with an update scheme (‘synchronous’ or ‘asynchronous’). If all nodes are updated simultaneously, the update scheme is referred to as synchronous [35]; otherwise it is said to be asynchronous [12]. If the *i*^th^ node is regulated by *k* inputs, then its state at the next time step is given by 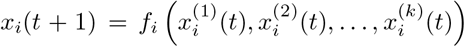, where *f*_*i*_ denotes the BF associated with node *x*_*i*_, and 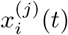 represents the *j*^th^ input to this BF at time *t*. Under synchronous dynamics, for which the BFs are applied at all nodes simultaneously, the state of the network goes from **X**(*t*) to **X**(*t* + 1) in one time step. Application of the BFs to all possible states defines the so called state transition graph (STG) wherein each node represents a distinct network state, and a directed edge indicates the transition between the associated states according to the BFs. Owing to the finite size of the state space of a BN, which consists of 2^*N*^ different states, repeated application of the rules on any network state leads to recurrent behavior, i.e., the system revisits states that it has already encountered. Under synchronous dynamics, the system enters attractors that are either fixed points, which remain unchanged under subsequent updates (i.e., **X**(*t* + 1) = **X**(*t*)), or limit cycles, i.e., the network perpetually cycles through a finite sequence of states.

### 2.2. Threshold functions and their subtypes

Threshold functions (ThFs) play a very prominent role in modeling regulation in both artificial neural networks and gene regulatory networks (GRNs) [16, 19, 21–23, 41–43]. In such a framework, the state of a given node (e.g., a gene) is determined by the collective effect of its activatory and inhibitory regulators [16, 44]. Let, *x*_*i*_ denotes the target node, and assume it receives signals from *k* regulators *x*_1_, *x*_2_, …, *x*_*k*_. A ThF assigned at the node *x*_*i*_ can be expressed as

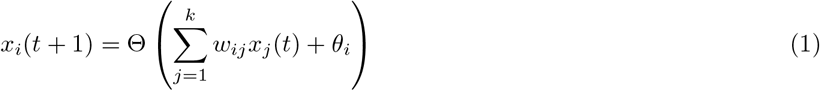

where *w*_*ij*_ *∈*ℝ is the weight (i.e., regulatory strength) of the directed edge from *x*_*j*_ to *x*_*i*_, and *θ*_*i*_ *∈*ℝ is the threshold for node *x*_*i*_. Θ(*s*) is an indicator function, returning ‘on’ (1) if *s >* 0, ‘off’ (0 or *−* 1) if *s <* 0, and a predefined value when *s* = 0.

The class of functions given by Eq. 1 is also known as linearly separable Boolean functions (LSBFs) [45, 46] when using a geometric point of view. A *k*-input BF may be represented as a *k*-dimensional hypercube where each of the 2^*k*^ vertices represents one input combination and is assigned with one of the two possible outcomes ‘on’ or ‘off’. LSBFs (i.e., ThFs) constitute a specialized subclass of BFs whose hypercube vertices can be partitioned by a single hyperplane into ‘On’ vertices on one side and ‘Off’ vertices on the other. Furthermore every ThF is unate, i.e., is monotonic in each of its inputs. Thus ThFs form a subclass of the family of unate functions (UFs) (see SI Text, Section 1) [47, 48]. Up to *k* = 3, the classes of UFs and ThFs are identical. For *k≥* 4, ThFs form a proper subset of UFs.

The exact numbers of ThFs are known for *k ≤* 9 [49]. However, we are really only interested in *effective* ThFs (EThFs, see SI text, Section 1) [36]. The exact numbers of EThFs have been published only for *k ≤* 8 [17]. However, the counts of EThFs can be easily computed using a recursion for any *k* up to 9. Let |ThF_*k*_| denote the total number of ThFs on *k* inputs, and let 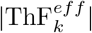 denote the the number of EThFs of *k* inputs. Every ThF in ThF_*k*_ is either an EThF or obtained by extending an EThF on *i < k* inputs to a *k* input BF by adding (*k−i*) dummy inputs. Given that there are exactly two constant BFs for any *k* (constant BFs are also ThF), and that each effective function (EF) on *i* inputs (see SI text, Section 1) [36] can be embedded into a *k* input BF in 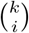 ways, we have:

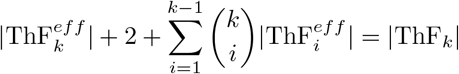

Rearranging,

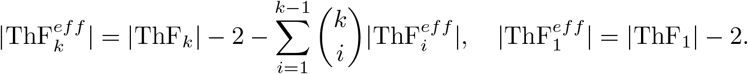

This recursion allowed us to calculate the number of EThFs up to *k* = 9; those values are provided in the SI Table S1.

It has been shown that the nested canalyzing functions (NCFs) [40, 50, 51] form a very specific subtype of ThFs. To be more specific, the NCF class is the intersection of the read-once functions (RoFs) (see SI text, Section 1) [52] and ThFs [53]. This means that suitable choices of *w*_*ij*_ and *θ*_*i*_ in Eq. 1 can yield any NCF. A *k*-input BF *f* is called NCF if there exists a permutation *π* of {1, 2, …, *k*}, canalyzing input values *a*_1_, *a*_2_, …, *a*_*k*_ *∈* {0, 1}, and canalyzed output values *b*_1_, *b*_2_, …, *b*_*k*_ *∈* {0, 1} such that:

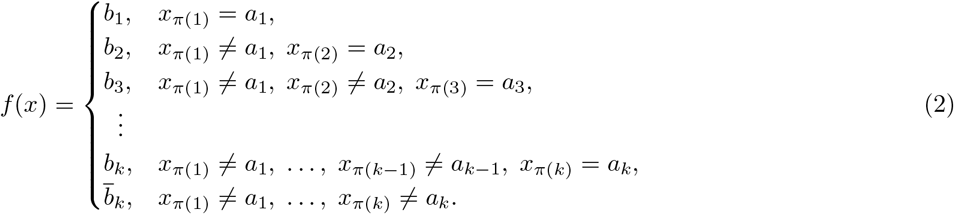

Alternatively, an NCF *f* with *k* inputs can be written as

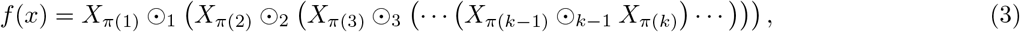

where *π* is a permutation of {1, 2, …, *k*}, ⊙_*i*_ ∈ {∧, ∨} for each *i*, and 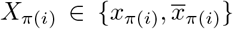. It has been shown in previous research that in spite of being a minuscule subspace of all BFs, NCFs are highly enriched in empirical datasets of regulatory logic rules [37].

In most experimental studies only the signs of the interactions (i.e., activation or inhibition) are reliably measured, while the precise strength of that regulation remains uncertain. To proceed with modeling the dynamics of GRNs, modelers often adopt the simplest possible ‘unit-weight’ assumption: each activatory or inhibitory edge from node *x*_*j*_ to *x*_*i*_ is assigned as *w*_*ij*_ = +1 or *−* 1, respectively while *w*_*ij*_ = 0 if no edge exists. Thresholds are likewise set to 0 (*θ*_*i*_ = 0), so that the update of node *x*_*i*_ depends only on the algebraic sum of its incoming signals and (potentially) the value of *x*_*i*_ itself. If that sum is positive, the node switches ‘on’; if it is negative, it switches ‘off’; and if it is exactly zero, the node retains its previous state. To date, modelers have typically employed one of the two binary conventions for the node states when using this simple threshold-based rules:

(a) Each *x*_*i*_ takes values in {*−*1, +1} which we refer as *Ising majority rule* or IMR [27, 29].

(b) Each *x*_*i*_ takes values in {0, 1} which we refer as *Bit-based majority rule* or BMR [31, 32]. More formally:

- **IMR update:** Suppose in a network with *N* nodes, the node *x*_*i*_ is regulated under IMR by *k* regulators *x*_*j*_ for *j* = 1, 2, …, *k*. Then its state at time *t* + 1 is given by:

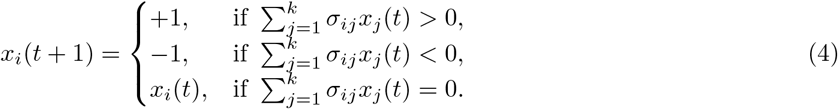
- **BMR update:** Suppose in a network with *N* nodes, the node *x*_*i*_ is regulated under BMR by *k* regulators *x*_*j*_ for *j* = 1, 2, …, *k*. Then its state at time *t* + 1 is given by:

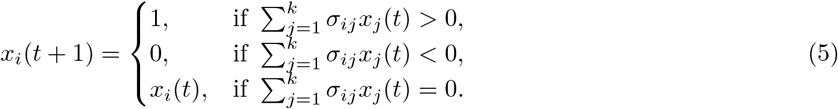

In IMR *x*_*i*_ *∈* {*−*1, +1} and in BMR *x*_*i*_ *∈* {0, 1} for each *i ∈* {1, 2, …, *N*} and in both formalisms one has

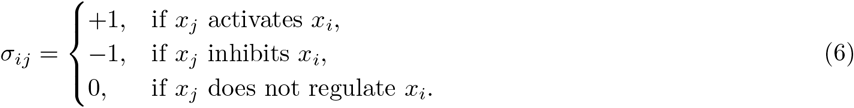

Note that, in the case of BMR, if an input is off, it does not affect the output, irrespective of whether the input is activatory or inhibitory. However, this is not true in case of an IMR. We refer to both of these variants of simple threshold-based rules collectively as *Threshold Majority Rules* or TMRs.

### 2.3. Empirical datasets for enrichment analysis of threshold functions and their subtypes

We consider three empirical datatsets of regulatory Boolean logic rules for the quantification and enrichment analysis of EThFs and of their different subtypes. Similar datasets were utilized in our previous work [54]. All datasets are publicly available via the GitHub repository: https://github.com/asamallab/GenChF. Characteristics of these datasets are as follows:

- **BBM benchmark dataset:** This dataset was obtained from the manually reconstructed models compiled by Pastva *et al*.[55] by first extracting 5990 BFs with a maximum of 10 inputs as described in [54]. For the analyses presented in this study, we further ensured that the BFs were EF. If a BF included any ineffective inputs, we removed those inputs to obtain the effective form. We then restricted our study to BFs with at most 9 effective inputs, since the theoretical count of ThFs is currently available only up to 9 inputs [49]. After applying these criteria, we obtained a final set of 5953 BFs. The distribution of these BFs based on input size and their classification into EThFs and its various subtypes is presented in SI Table S2.
- **MCBF dataset:** This dataset is based on a previously published study [37]. Starting with that earlier case, we first converted any ineffective BFs into their effective forms by removing ineffective inputs. We then kept the BFs with at most 9 inputs. The final dataset for our analyses comprises 2670 BFs. The distribution of these BFs based on input size and their classification into EThFs and its various subtypes is presented in SI Table S3.
- **Harris dataset:** This dataset was published in [40, 56]. It contains 149 BFs, each with at most 5 inputs. Therefore, all BFs were considered exhaustively for our analysis. The distribution of these BFs based on input size and their classification into EThFs and its various subtypes is presented in SI Table S4.

### 2.4. Relative enrichment and statistical significance

To test whether EThFs within all BFs or different subtypes of EThFs within EThFs (e.g., NCF, IMR, BMR) are overrepresented or underrepresented in an empirical datatset, we adopt a relative enrichment framework as presented in Subbaroyan *et al*. [37]. *Let T* denotes a class of BF (e.g., *T* = EThF) and *T*_*s*_ *⊂ T* one of its subtypes (e.g., *T*_*s*_ = IMR). Working at a fixed number of effective inputs *k*, let us define:

- *f*_*s*,1_ = Fraction of *k*-input BFs in the dataset that lie in *T*_*s*_.
- *f*_1_ = Fraction of *k*-input BFs in the dataset that lie in *T*.
- *f*_*s*,0_ = Fraction of all possible *k*-input BFs that lie in *T*_*s*_.
- *f*_0_ = Fraction of all possible *k*-input BFs that lie in *T*.

The relative enrichment of *T*_*s*_ within *T* is then

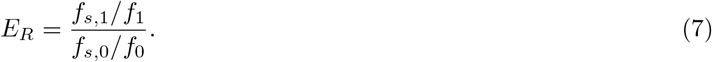

An *E*_*R*_ value *>* 1 indicates that the subtype *T*_*s*_ is proportionally more common within *T* in the empirical dataset than it is in the universe of all *k*-input BFs. On the other hand *E*_*R*_ value *<* 1 is indication of underrepresentation of that subtype *T*_*s*_ within *T* in the empirical dataset. We also accompany each *E*_*R*_ estimate with a formal significance test, as described below.

### Assessing statistical significance

Under the null hypothesis *H*_0_, we assume that although the empirical dataset may be enriched overall for BFs in *T*, the selection among members of *T* is random, that is the probability of elements in *T*_*s*_ is not more than other elements in *T*. Equivalently, given that *M*_*T*_ of the empirical dataset’s *M* total BFs lie in *T*, each of those *M*_*T*_ slots is equally likely to be occupied by any member of *T*, irrespective of whether it belongs or not to *T*_*s*_. Let, *f*_*R*_ = | *T*_*s*_ | */* | *T*| be the background probability that a randomly chosen BF from *T* belongs to *T*_*s*_. Then, under *H*_0_, the number *m* of *T*_*s*_ members observed among the *M*_*T*_ selected BFs follows a binomial distribution:

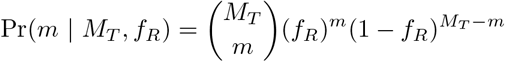

The desired one-sided *p*-value for the observed count *m*_*obs*_ is:

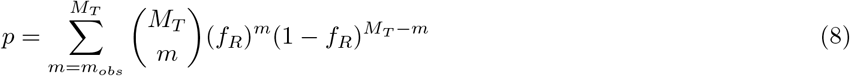

A small *p*-value indicates that the observed enrichment of *T*_*s*_ within *T* in the empirical dataset is unlikely under the statistics given by *H*_0_. Conversely, if *T*_*s*_ is not enriched within *T*, or is even underrepresented within *T*, the calculated *p*-value will no longer be small or it may even approach 1, signaling that the observed frequency is consistent with *H*_0_ or is associated with depletion.

### 2.5. Complexity Measures

A range of complexity metrics for BFs have been explored in the computer science literature [57, 58]. In this study, we focus on two of them: Boolean complexity [38] and average sensitivity [39].

#### Boolean complexity

One way to quantify a BF’s succinctness is via its so-called Boolean complexity. This measure, formulated by Feldman [38], reflects the minimal descriptive length of a BF by counting the number of literals in its shortest equivalent form. While canonical forms like conjunctive normal form (CNF) or disjunctive normal form (DNF) uniquely represent a BF, they often contain redundant terms that can be eliminated using Boolean algebra. A minimal formula is obtained by systematically applying these simplifications to collapse multiple product or sum terms into a more compact form. For instance, the 3-input BF

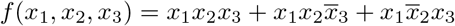

initially appears with nine literals in its canonical DNF form. However, applying Boolean algebra, one can derive a minimal equivalent expression

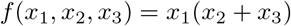

which uses only three literals. Determining an absolute minimal formula is a 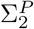 -complete problem [59, 60], so in practice one employs a heuristic logic-synthesis approach. As described in Subbaroyan *et al*. [37], in our pipeline, each BF is first converted into four candidate representations: full DNF, full CNF, Quine-McCluskey minimized DNF and Quine-McCluskey minimized CNF [61, 62]. We then pass each candidate to the “ABC” logic-synthesis tool [63] which perform further algebraic optimization. Among the four outputs, the one with the fewest literal counts serves as a proxy for the BF’s true Boolean complexity.

#### Average sensitivity

The average sensitivity [39] of a BF quantifies its propensity to change output under single-bit perturbations of its inputs. For a *k*-input BF *f*, it is defined as

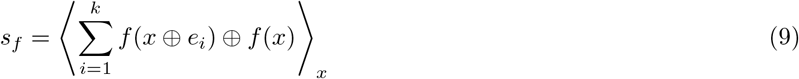

where *⊕* denotes the logical XOR operator, *e*_*i*_ *∈* {0, 1}^*k*^ is the unit vector with a 1 in the *i*^th^ position and 0 elsewhere, and *⟨·⟩*_*x*_ denotes the average over all 2^*k*^ binary inputs *x*.

### 2.6. Imposing structural and dynamical constraints on biological models

We assembled a collection of manually curated BN models to evaluate whether the simple TMRs suffice to reproduce key dynamical behaviors. Our procedure comprises a rigorous multi-step selection as summarized in SI Fig. S1. Starting with the Biodivine Boolean Models (BBM) benchmark dataset [55], which currently contains 245 models drawn from diverse databases and literature, we first discard models that are not manually assigned or that employed multi-level (non-Boolean) variables. From the remaining set, we then removed models lacking a peer-reviewed reference. To ensure compatibility with exhaustive attractor and state transition enumeration using the R package ‘BoolNet’ [64], we retained only those models with fewer than 30 nodes. To have consistency with other analyses in this work, we also removed the models with a maximum in-degree of 10 or more. Models that incorporated any ineffective BF were subsequently discarded. At this point, each remaining model is then cross-checked against the corresponding publication to confirm that the underlying GRN topology and the sign (i.e., activatory or inhibitory) of each regulatory interaction were reported and consistent with the model. This is critical for the subsequent replacement of each original BF with the corresponding TMRs. Finally, we retained only those models for which the biological attractors were explicitly described in the publication and were recovered by simulating the model. This filtration pipeline yielded 24 models for further analysis. Of these, 20 had all their BFs belonging to the NCF class, while each of the remaining 4 models had at least one non-NCF BF.

### 2.7. An attractor-based scoring framework to compare a published model to an alternative model

We would like to evaluate how well an alternative model formed by replacing each BF of the published model by a different rule (specifically, by the corresponding TMR variants) reproduces the given biological outcomes. In particular, we ask: how accurately does the alternative model recover the literature-reported biological attractors, and to what extent does it preserve the associated basin size distributions? To address this, we propose two metrics:

#### (a) Attractor Recovery Score and (b) Distance of basin fraction distribution from the gold standard

For any given model in our empirical dataset, let *A*_*bio*_, *A*_*model*_ and *A*_*alt*_ denote the set of attractors considered as biologically relevant in the publication (hereafter, referred to as the biological attractors), the set of actual attractors produced by the published model, and the set of attractors of the alternative model, respectively. By construction (see Section 2.6), the considered published models always recover the biological attractors, so that *A*_*bio*_ *⊆ A*_*model*_. To quantify attractor recovery performance, we introduce the attractor recovery score (ARS), defined as the ratio of two Jaccard indices. First, let

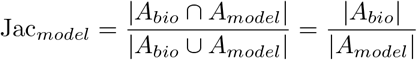

where the equality follows from *A*_*bio*_ *⊆ A*_*model*_. Next define,

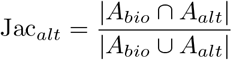

The attractor recovery score for the alternative model is then

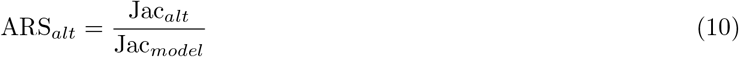

A score below one indicates degraded performance relative to the published model, either because the alternative model fails to recover all biological attractors or because it generates many spurious attractors or both. In the extreme case, a score of zero or close to zero indicates almost no recovery of the biological attractors or such an abundance of attractors that any overlap with biological attractors is random. A score of exactly one signifies that the alternative model matches the published model’s performance. Finally, a score above one denotes improved performance, signifying that the alternative model not only recovers every biological attractor but does so with fewer spurious attractors than the published model.

In synchronous BN dynamics, the entire state space partitions into disjoint basins of attraction (BoA), each associated with a particular attractor. The BoA for a given attractor comprises all initial states from which the system eventually converges to that attractor. Thus, by normalizing each basin’s size by the total number of states, one obtains a probability distribution over attractors, where each basin fraction corresponds to the likelihood of the system converging to that attractor from a randomly chosen initial state. To quantify how well a model reproduces the expected basin size distribution, we employ the Jensen-Shannon (JS) distance [65, 66]. In an ideal scenario, we anticipate that a good model would recover only the biological attractors, with no spurious ones. However, the published models quite often exhibit additional artefactual attractors, i.e. *A*_*bio*_ is a proper subset of *A*_*model*_. Since, the true “gold standard” basin fractions of the biological attractors are unknown a priori, we take the published model’s basin fraction for the attractors in *A*_*bio*_, renormalize them over only the biological attractors, and use this renormalized distribution for our reference “gold standard”. Formally, let

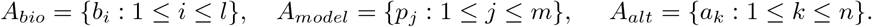

Now, clearly *l ≤ m* as *A*_*bio*_ *⊆ A*_*model*_. Let 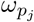 denote the published model’s basin fraction of attractor *p*_*j*_ for all 1 *≤ j ≤ m*. Without loss of generality, assume *b*_*i*_ = *p*_*i*_ for 1*≤ i ≤ l*. The renormalized (“gold-standard”) basin fraction for each biological attractor is then given by:

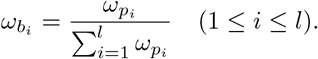

We now form two probability vectors of length *m*:

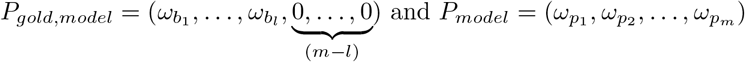

where the trailing zeros in *P*_*gold,model*_ accounts for the (*m − l*) artefactual attractors of the published model. The JS distance *D*_*model*_ between these two vectors quantifies the published model’s deviation from the ideal expectation. *D*_*model*_ = 0 arises only when the published model does not generate any artefactual attractors.

For the alternative model, let *r* = |*A*_*bio*_ *∩A*_*alt*_| and, again without loss of generality, assume *b*_*i*_ = *a*_*i*_ for 1 *≤ i ≤ r*. If *l > r*, then the alternative model fails to recover (*l − r*) biological attractors and if *n > r* then the alternative model generates (*n − r*) spurious attractors. Therefore, we construct two probability vectors of length *l* + *n − r*:

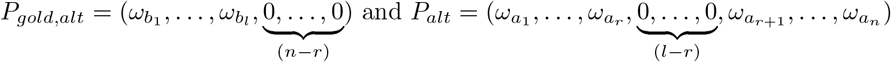

where 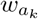 denotes the alternative model’s basin fraction for attractor *a*_*k*_, 1 *≤ k ≤ n*. The JS distance *D*_*alt*_ between these vectors is the alternative model’s deviation from the ideal expectation. By comparing *D*_*model*_ and *D*_*alt*_, both of which lie in [0, 1], we can assess relative performance of the alternative model. A lower distance corresponds to better performance.

### 2.8. Stability of Boolean network dynamics

#### Stability measures based on damage spreading

For assessing the dynamical stability of BNs [67], a common approach is to determine, how a localized perturbation such as a single-bit flip propagates over time. This approach is formalized by the Derrida map [68], which relates the average Hamming distance or ‘damage’ between two trajectories at successive steps. If small perturbation tends to decay then the network dynamics is deemed stable, and if they tend to amplify then the dynamics exhibits chaotic behavior. We employ three different metrics [69] based on damage spreading: the Derrida coefficient (*δ*), the final Hamming distance (*h*^*∞*^) and the fragility (*ϕ*). All averages are taken over all possible initial states and over single-bit perturbations.

The Derrida coefficient (*δ*) is defined as the average Hamming distance between two trajectories at time (*t*+1), given that their initial Hamming distance at time *t* was exactly 1. It reflects how a small initial perturbation propagates in the network after a single update. *δ <* 1 indicates that perturbations tend to die out (ordered regime), while *δ >* 1 indicates that perturbations spread to a finite fraction of nodes (chaotic regime).

The measure final Hamming distance (*h*^*∞*^) measures the long-term average separation between unperturbed and perturbed trajectories. Formally,

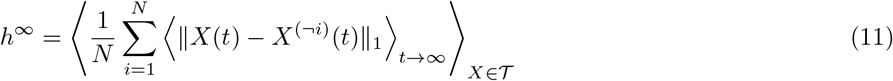

where *X*(*t*) is the unperturbed network state at time *t* and *X*^(*¬i*)^(*t*) denotes the state evolved from the initial condition where a single-bit perturbation was appied to the *i*^th^ variable. ∥ · ∥_1_ denotes the Hamming distance, ⟨·⟩_*t→∞*_ is the long-time average, and ⟨·⟩ _*X∈T*_ denotes the average over all initial states.

The fact that *h*^*∞*^ can be affected by phase shifts in cyclic attractors motivated the introduction of the so called fragility (*ϕ*). That measure averages the time series first and then computes the distance, removing any sensitivity to phase differences, and thus, provides a robust measure of long-term divergence. Mathematically,

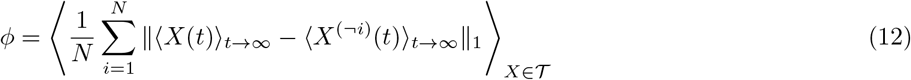

Similar to *δ*, the values *h*^*∞*^ = 1 and *ϕ* = 1 are taken as critical thresholds: values below this boundary indicate robust or ordered dynamics, while values above suggest chaotic behavior.

#### Measures based on attractors and on the state transition graph to characterize the dynamics

In addition to the three previous quantities based on damage spreading, we use stability measures associated with the attractors or the state transition graph (STG) as follows:

i. Total number of attractors: This refers to the combined count of all fixed point and cyclic attractors identified in the system. BNs with high stability tend to have fewer attractors [67].
ii. *G*-density: This quantity is the fraction of garden-of-Eden (*GoE*) states in the STG, i.e., the fraction of states with in-degree 0 [70, 71]. A higher *G*-density corresponds to more irreversible and thus more robust dynamics.
iii. Average convergence rate of the trajectories originating at *GoE* states (*λ*_*GoE*_): This is the inverse of the average of the lengths of transients (steps to the attractor) originating at *GoE* states [71]. A higher convergence rate is expected for robust dynamics.

### 2.9. Ensembles of random Boolean networks with topological constraints and Boolean function classes as subtypes of threshold functions

Previous studies on random Boolean networks (RBNs) adopted specific choices to define the architecture, commonly referred to as network topology. Two widely-used topological ensembles are as follows. In the first, directed edges are assigned randomly between node pairs, maintaining a fixed average number of edges *λ* per node. This setup corresponds to a directed version of the Erdös-Rényi [72] network architecture. We refer to it as the P-P ensemble as both the in-degree (*k*_in_) and out-degree (*k*_out_) distributions converge to a Poisson distribution with mean *λ* in the large network limit. In the second ensemble, both the in-degree and out-degree of each node are fixed to a constant value, resulting in what is known as a *k*-regular graph ensemble, hereafter referred to as the R–R ensemble [73].

#### Simulation setup

We generate and analyze ensembles of RBNs under two distinct topological configurations: P-P and R-R (as described above). The analyses are performed as follows: (a) We consider networks with *N* = 12 and *N* = 16 nodes. (b) The average in-degree (*λ*) or fixed in-degree (*k*_*in*_) considered takes values 2, 3, 4 and 5. (c) For networks with *N* = 12, we generate 10000 RBNs per setting, while for *N* = 16, we generate 1000 RBNs per setting. (d) In each case, for every RBN, we generate three different models: one where all BFs are sign conforming NCFs (scNCFs), one where all are set to BMRs, and one where all are set to IMRs. (e) With this setup, now we compute different measures for each Boolean model as described in Section 2.8.

## 3. RESULTS

Due to paucity of information on activation thresholds or regulatory weights, it seems natural to resort to threshold majority rules (TMRs) as they do not require such information. TMRs thus offer an appealingly simple and uniform framework to model the dynamics of complex biological networks. However, it is important to ask whether this simplicity comes at the cost of biological realism. The broader question we address in this study is whether TMRs are appropriate for modeling biological gene regulatory networks (GRNs). To be more explicit, does their use lead to behavior consistent with the properties of biological GRNs as found in reconstructed networks? To investigate this, we undertake a systematic and comprehensive analysis of both variants of TMRs: (a) Ising majority rules (IMRs) and (b) Bit-based majority rules (BMRs). Our results are organized across five subsections. The first three focus on function-level analyses, while the final two shift to probing the dynamics at the network level.

In the first subsection, we begin by identifying the Boolean functions (BFs) that are equivalent to TMRs in the space of all BFs. We examine whether these rules exhibit properties such as odd bias and canalyzation, which are commonly observed in empirically curated BFs [37]. In the second subsection, we examine the empirical relevance of TMRs by assessing their presence in empirical datasets and determining whether they are enriched or underrepresented relative to theoretical expectations. In the third subsection, we turn to their regulatory complexity. While TMRs are often chosen for ease of implementation, we explore whether they are likewise simple in terms of their intrinsic logic. In the fourth subsection, we shift our focus to network dynamics. Since attractors correspond to key biological outcomes and a good model must recover both these attractors and their basin size distributions, we evaluate whether TMRs fulfill this requirement. Finally, in the fifth subsection, we perform an analysis of the stability of the dynamics by measuring a number of quantitative metrics to determine whether networks constructed using TMRs exhibit ordered behavior or lean more toward chaotic dynamics.

Taken together, these investigations aim to determine whether the simplicity of TMRs justifies their continued use as a modeling framework, or whether they warrant a more critical stance.

### 3.1. Representation of threshold majority rules as Boolean functions and their properties

To make a meaningful comparison and to understand different properties of TMRs (see Section 2.2), it is essential to interpret TMRs in a common framework with nested canalyzing functions (NCFs) or any other BF classes. Specifically, we want to explore the full range of input-output behavior, i.e., identify the BFs that correspond to IMRs and BMRs under different sign combination of the regulators and their properties (See Fig. 1).

**FIG. 1.**
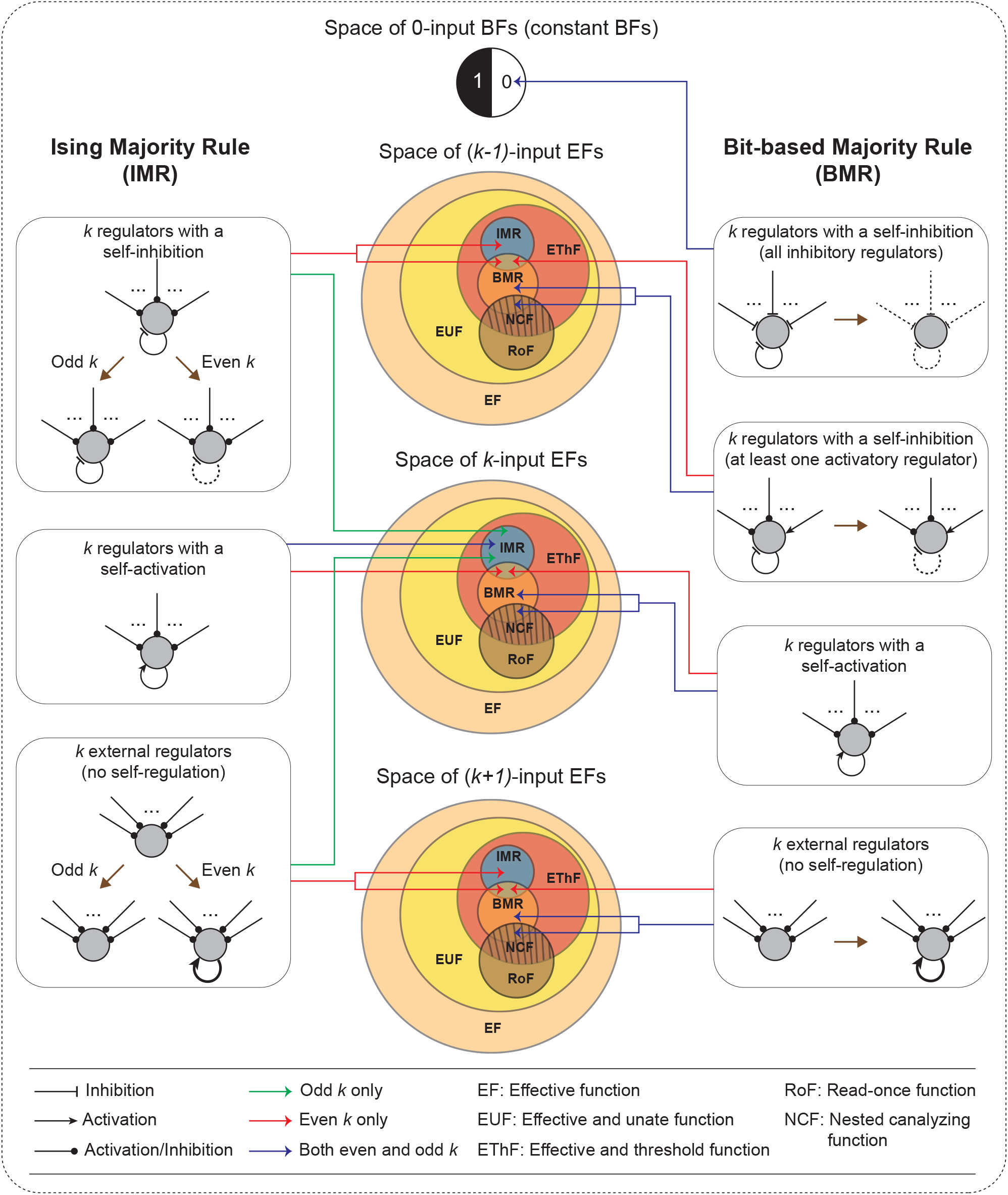
A schematic depiction of the mapping of TMRs to the space of EFs for *k >* 2. This schematic illustrates the correspondence between the TMRs and the subsets of EFs, across all possible interaction topologies of regulators at a target node. The mapping from the TMR formulation to the usual BF formulation keeping only effective interactions is indicated by the brown arrows. The target node is depicted via a grey circle. The three major Venn diagrams represent the space of EFs having (*k −* 1), *k* and (*k* + 1) inputs, respectively (note that areas of the subspaces within each Venn diagrams are not to scale). Dashed inhibitory edges (either self-regulatory or external) indicate their ineffectiveness under specific TMR conditions (topology, parity, signs). The absence of a self-regulation can lead to added self-activations as shown using thick black self-loop arrows. Inversely, even though a TMR may include an explicit self-inhibition, the corresponding EF may have no self-interaction. Colored lines connect each topology box to the corresponding regions in the Venn diagrams. Red lines indicate mappings for even *k*, green lines for odd *k* and blue lines for both even and odd values of *k*.

We first begin with the IMRs. Assume that a node *x*_*i*_ is governed by an IMR and is regulated by *k external* regulators *x*_*j*_, that is external to *x*_*i*_ so that there is no self-regulation (sometimes referred to as self-loop or autoregulation or self-interaction). Each *x*_*j*_ to *x*_*i*_ influence has a predefined sign *σ*_*ij*_, representing either activation or inhibition, where 1 *≤ j ≤ k*. Now, when *k* is odd, the signed sum 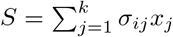 can never be zero given that each *x*_*j*_ *∈* {*−*1, +1}. In this scenario, the output is determined entirely by the inputs from its regulators and does not depend on its own state. As a result, the corresponding BF belongs to the space of all *k*-input effective functions (EFs) (see Fig. 1 and SI text, Section 1) [36]. However, the situation changes when *k* is even, since then the sum *S* will be zero for certain input combinations. When this happens, the output of the rule depends on the current state of *x*_*i*_ as well, despite the fact that *x*_*i*_ is not structurally self-regulating. In other words, IMRs introduce an (implicit) self-activation, and consequently, the corresponding BF must now be interpreted as a (*k* + 1)-input EF (see Fig. 1). Now consider the case when the set of *k* regulators of the node *x*_*i*_ governed by IMR includes *x*_*i*_ as well. In principle, such a system corresponds to a *k*-input BF. However, not every such BF remains EF. Indeed, when *k* is even and the self-regulation is inhibitory, the BF becomes ineffective: the self-regulation (and thus *x*_*i*_) does not influence the output, while other (*k−*1) inputs remain effective (see Fig. 1). Furthermore, for *k* = 2, even when the self-regulation is activatory, the BF is not effective. In more detail, suppose *x*_2_ is regulated by both *x*_1_ and *x*_2_. Under the four possible sign configurations for (*x*_1_, *x*_2_): ++, *−*+, *−*+ and *−−*, the rule reduces to the single input EFs: *x*_2_, *x*_1_, *x*_2_ and *x*_1_, respectively. Thus, there is no two-input EF which corresponds to an IMR. However, when *k* is odd, any IMR still corresponds to a *k*-input EF even if there is a self-regulation because the signed sum of all inputs can never equal zero, just as in the case with no self-regulation.

We can further characterize the bias (the number of input combinations yielding an ‘on’ output) of an EF *f* which is equal to an IMR with *k* regulators: (a) for odd *k*, the bias is 2^*k−*1^; (b) for *k* = 2, the bias is 1; (c) for even *k ≥* 4, an external set of regulators produces a bias of 2^*k*^; (d) for even *k≥* 4, a set of regulators that includes an activatory self-regulation produces a bias of 2^*k−*1^; and (e) for even *k≥* 4, a set of regulators that includes an inhibitory self-regulation produces a bias of 2^*k−*2^ (see SI text, Properties 2.1 and 2.2). This pattern implies that any EF which is equal to an IMR with more than two regulators (with or without a self-regulation) and any two-regulator

IMR without a self-regulation will always have an even bias. In an empirical survey of BFs extracted from published Boolean models, some of us previously showed the predominance of odd-biased BFs [37]. Moreover, no BF equal to an IMR with more than two regulators can be canalyzing (see SI text, Section 2.1 and Property 2.3), standing at odds with both Kauffman’s theoretical framework of canalyzation [35] and the prevalence of canalyzing functions (CFs) in empirical datasets [37].

Compared to IMRs, BMRs operate on a fundamentally different principle. Given a BMR, the sign of an interaction plays no role if the corresponding input is inactive; in a voting analogy, that input provides only a blank vote, whereas in IMRs all inputs vote regardless of whether they are active or inactive. Clearly, the output of a BMR is determined solely by comparing the total regulatory strength of the active activators and of the active inhibitors. Consider first the case where a target node *x*_*i*_ has *k* external regulators *x*_*j*_. There is always at least one input configuration, specifically the one where all *x*_*j*_ = 0, for which the signed sum is 0; for that input, the output depends on the value of *x*_*i*_. Thus a BMR with *k* external inputs corresponds in fact to a (*k* + 1)-input EF, whether *k* is even or odd (see Fig. 1).

The situation becomes more nuanced if *x*_*i*_ is one of the explicit regulators itself, i.e., if a self-regulation is present (see SI text, Property 2.4). For *k >* 1, if the self-regulation is inhibitory and not all other regulators are also inhibitory, the self-regulation becomes ineffective and the BMR corresponds in fact to a (*k−* 1)-input EF. Interestingly, for *k >* 0, if all *k* regulators including the self-regulation are inhibitory, the rule collapses to a constant zero BF which clearly has no effective inputs (see Fig. 1). On the other hand, if the self-regulation is activatory, the BMR corresponds to a *k*-input EF for all *k >* 2 (see Fig. 1). However, for *k* = 1 and 2, it is a *k*-input EF only when all regulators are activatory.

Unlike the relatively straightforward bias properties of IMRs, the bias of BMRs also depends on the regulatory sign pattern via the number of activators and inhibitors. For a BMR with *k* external regulators, *a* of which are activators, the bias of the equivalent BF is given by (see SI text, Property 2.5):

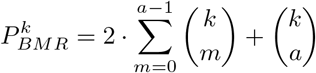

revealing that both even- and odd-biased EFs can arise depending on the values of *a* and *k*. In fact, one can show that odd bias arises precisely when 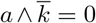, where *∧* denotes the bitwise AND operation on the binary representations of *a* and *k* (see SI text, Property 2.5, Corollary 2.1). Moreover, whenever *k* is a Mersenne number (one less than a power of two), all corresponding BMRs have an odd bias (see SI text, Property 2.5, Corollary 2.1). When the self-regulation is present, then the bias of the *k*-input BF equivalent to the given BMR is (see SI text, Property 2.6):

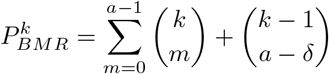

where *δ* = 0 for an activatory self-regulation and *δ* = 1 for an inhibitory one. Note that if the self-regulation is inhibitory or *k* = 2 and the regulators are not both activatory, then the BF is not effective and consequently for the reduced equivalent EF, the bias is halved, equal to 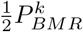.

Unlike IMRs, BMRs can be canalyzing if several conditions are met. A BMR over *k* external regulators is canalyzing if and only if either all regulators are activators or all are inhibitors (see SI text, Property 2.7). Moreover, such BFs are not only canalyzing but also NCFs (see SI text, Property 2.5, Corollary 2.1), specifically, they correspond to OR and AND type rules, respectively. When the set of regulators also include the target node, then the BF is canalyzing if at most one regulator is activatory or at most one is inhibitory (see SI text, Property 2.8).

Despite their many contrasting features, IMR and BMR can occasionally correspond to the same BF. To make this comparison, we treat BFs as equal if their truth table outputs match when we identify the values ‘-1’ (in IMR) and ‘0’ (in BMR) as the same logical ‘off’ state and ‘+1’ as the logical ‘on’ state. Under this correspondence, let *f*_1_ and *f*_2_ denote the BFs equal to an IMR and a BMR respectively, each with *k* regulators and the same sign configuration of the regulators. Then *f*_1_ and *f*_2_ are equal only if: (a) *k* = 1 and there is a self-activation, or (b) *k* is even and the number of activators equals the number of inhibitors (see SI text, Property 2.9).

These properties illustrate that in many cases the BFs corresponding to TMRs lack important logic features present in NCFs, and thus, lack features which are typically present in biological GRNs. For instance, IMRs almost always produce even-biased and non-canalyzing rules. BMRs allow more variations but still offer very limited scope of canalyzation. Yet the most striking shortcoming of both IMRs and BMRs is their mishandling of self-regulation, specifically inhibitory self-regulations. IMRs ignore inhibitory self-regulations (that is self-regulations with negative weights) whenever the number of regulators is even, while BMRs discard any inhibitory self-regulations altogether irrespective of the parity of the number of regulators. This behavior is at odds with biological reality, where negative auto-regulation often plays a crucial role. For instance, studies of the *Escherichia coli* transcriptional regulatory network indicate that nearly 70% of the transcription factors are self-regulating with two-thirds of those being of negative sign [74, 75]. Thus, while TMRs offer a simple modeling framework, they generally fail to capture key regulatory properties found in real biological systems.

### 3.2. Threshold majority rules are underrepresented in empirical datatsets

With the mappings described in the subsection 3.1, we now test whether the BFs that are equal to certain TMRs are preponderant in empirical datasets of regulatory logic rules. To this end, we consider three different empirical datasets (see Section 2.3): (a) the BBM benchmark dataset, (b) the MCBF dataset, and (c) the Harris dataset. Just as for the BFs in each empirical dataset, we ensure that BFs that are equal to some IMR or BMR are effective. Specifically, for any BF which is ineffective, we remove the ineffective inputs to make the BF an EF. Subsequently, we categorize these BFs in different classes based on their number of effective regulators *k*. As discussed in Methods (see Section 2.2), the IMR, BMR and NCF classes each represent a specific subtype of ThFs. Among these, NCFs are always effective. Therefore, the NCFs and the effective BFs that are equal to some IMR or BMR constitute specific subtypes of EThFs.

#### Case of the BBM benchmark dataset

For the BBM benchmark dataset, we first compute the fraction of EThFs within all BFs for each input size *k* up to *k* = 9. We observe that EThFs are consistently enriched across all values of *k*, as shown in Fig. 2(a). In a previous work by some of us, it was shown that NCFs are significantly enriched in the empirical dataset of regulatory logic rules [37]. This leads us to compute the fraction of NCFs within the space of EThFs in the BBM benchmark dataset, across different values of *k*, as illustrated by the bars in Fig. 2(b). The corresponding statistical significance (see Section 2.4) values indicate that even within the space of EThFs, NCFs show a very strong relative enrichment for all values of *k >* 2. Note that, for *k* = 1 and 2, the theoretical spaces of NCFs and EThFs completely overlap. However, as *k* increases the proportion of NCFs within the theoretical space of EThFs progressively decreases and becomes minuscule for large *k* (see SI Table S1).

**FIG. 2.**
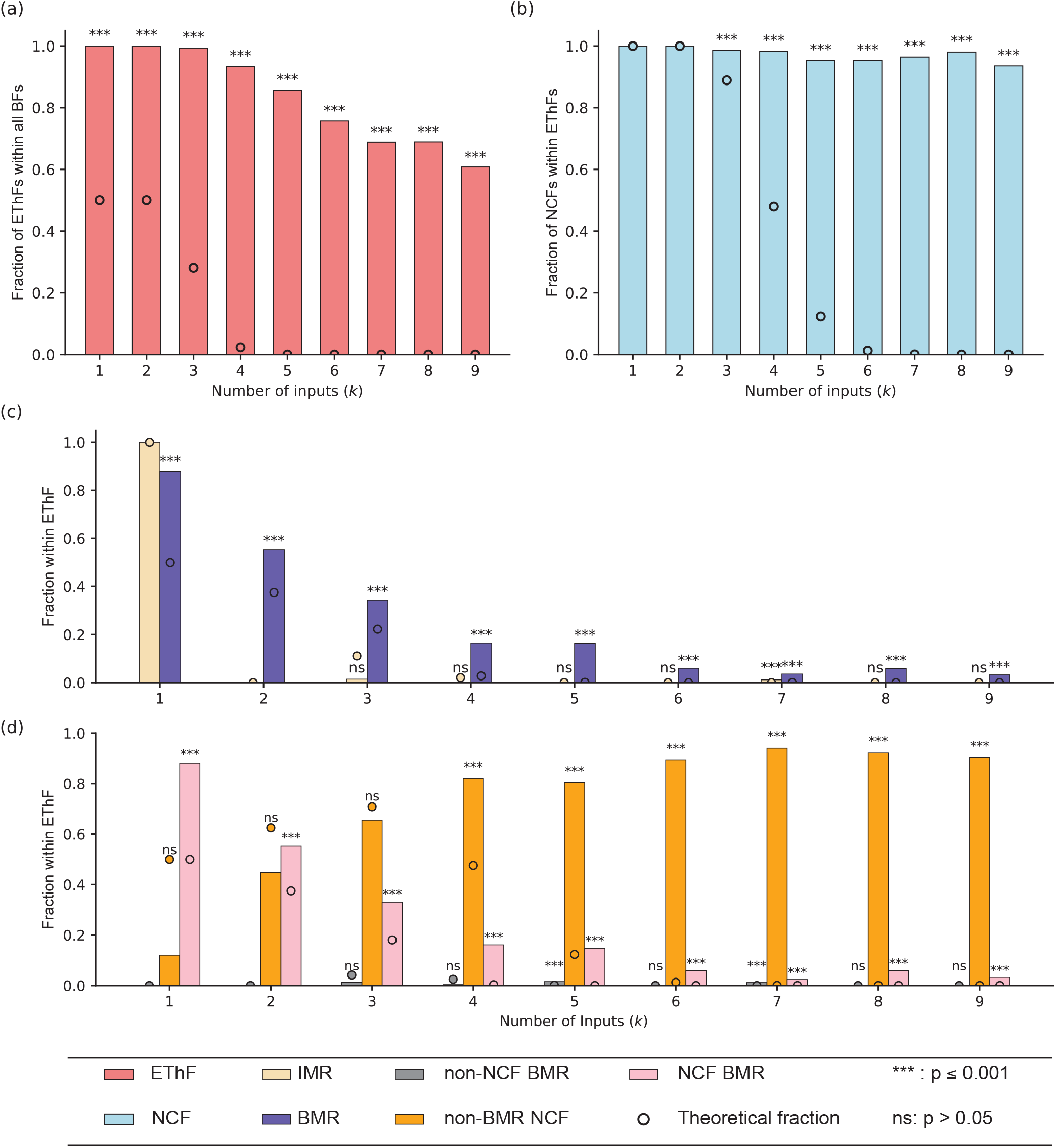
Fraction of various subtypes within an englobing type inside the BBM benchmark dataset. **(a)** The fraction of EThFs within all BFs for various number of inputs. **(b)** The fraction of NCFs within EThFs for various number of inputs. **(c)** The fraction of BFs that are equal to some IMR or BMR within the NCFs for various number of inputs. **(d)** The fractions of non-NCF BMRs, non-BMR NCFs and NCF BMRs within EThFs for various number of inputs. In all these plots, the theoretically expected values are represented by dots while the empirically obtained values from the BBM dataset are shown via the color bars.

We now turn to the same quantities for the other two subtypes of EThFs: IMR and BMR (see Fig. 2(c)). Note that for *k* = 1, the space of EThF, NCF and IMR completely overlap. Therefore, for *k* = 1, both the theoretical and empirical fractions of IMRs within EThFs are equal to 1. For *k* = 2, there are no EFs that are equal to any IMR, and hence, the theoretical fraction itself is 0. However, for *k >* 2, the BFs that are equal to some IMR form a proper subset of EThF, which is in fact disjoint from the NCFs (see SI text, Property 2.3). As shown in Fig. 2(c), for *k >* 2, the fraction of BFs equal to some IMR is generally 0 or very low within EThFs in the BBM benchmark dataset (see SI Table S2). In nearly all cases, these empirical fractions are lower than the corresponding theoretical expectations, indicating depletion rather than enrichment (see SI Table S5).

For all values of *k* up to 9, we observe that the BFs equal to BMRs are relatively enriched within the space of EThFs in the datatset (see Fig. 2(c) and SI Table S5). However, since for all *k*, there is a non-empty overlap between BMR and NCF (see SI text, Corollary 1.2), we must determine whether this enrichment is primarily due to BMRs that are also NCFs (NCF BMRs), or due to those that are not (non-NCF BMRs). To investigate this, we partition the set of BFs equal to BMRs into two categories for each *k*: (a) NCF BMRs (BFs that are both BMR and NCF) and (b) non-NCF BMRs (BFs that are equal to some BMR but are not NCF). For consistency, we also consider, for each *k*, the subset of NCFs that are not equal to any BMR, denoting them as non-BMR NCFs. For *k* = 1 and 2, all BMRs are also NCFs, so the set of non-NCF BMR is empty (see SI Table S1). However, for *k ≥* 3, the set of non-NCF BMR becomes non-empty. Given these definitions, we compute for each value of *k* the fraction of BFs belonging to these three categories within EThFs in the dataset along with their associated *p*-values. As can be inferred from Fig. 2(d), the observed enrichment of BMRs within EThFs is largely driven by the subset NCF BMR. The empirical fraction of non-NCF BMRs within EThFs is typically lower than the corresponding theoretical expectation. In contrast, non-BMR NCFs are highly enriched within EThFs for all *k >* 3, and this is statistically significant. Furthermore, a comparison of the associated *p*-values for the three categories reveals a consistent hierarchy for *k >* 3 which we denote as follows: non-BMR NCF *<* NCF BMR *<* non-NCF BMR (see SI Table S6).

#### Cases of the MCBF and Harris datasets

We performed the analogous analyses on the MCBF and Harris datasets and found results consistent with those observed for the BBM dataset (see SI Figures S2 and S3). In the MCBF dataset, the BFs equal to IMRs are not enriched for all *k >* 2. For *k >* 3, there are no BFs present in the MCBF dataset that are equal to IMRs (see SI Fig. S2, and SI Tables S3 and S7). Moreover, similar to what was found using the BBM dataset, the empirical fractions of non-NCF BMRs within the EThFs are generally low and statistically insignificant in the MCBF dataset (see SI Fig. S2). Here also, for *k >* 4, the associated *p*-values of the three classes exhibit the same consistent hierarchy: non-BMR NCF *<* NCF BMR *<* non-NCF BMR (see SI Table S8). We further repeated the analyses on an earlier dataset compiled by Harris *et al*. [40, 56], *and observed comparable results. In this dataset, the BFs have at most 5 inputs. For all values of k*, there are no non-NCF BFs that are equal to any IMR or BMR (see SI Fig. S3 and SI Table S4). See SI Tables S9 and S10 for the enrichment ratios and the corresponding *p*-values.

These findings show that IMRs and BMRs are not only underrepresented in the empirical datasets of biological regulatory logic rules but are in fact depleted compared to what would be expected by random selection. These empirical datasets compile the best available logic rules, each carefully assigned by experts around the world. While individual choices may reflect particular biases, these rules collectively can be considered the most reliable portrait of biological regulatory logic. From our analysis, it is clear that the TMRs fail to align with these curated logic rules. One may thus speculate that TMRs are not simply rare but might even be fundamentally incompatible with the logic underlying real biological networks.

### 3.3. Threshold majority rules exhibit higher averaged Boolean complexity and sensitivity than nested canalyzing functions

In a previous work by some of us [37], the low average values of sensitivity of the NCF set and of the Boolean complexity of its superset RoF were proposed as a plausible explanation for their noticeable enrichment in empirical datasets. That work also demonstrated that, for any given number of effective inputs *k*, RoFs and consequently NCFs attain the minimum possible Boolean complexity (see Section 2.5), which is equal to *k*. This result also directly follows from Eq. 3. Furthermore, in the context of the so called *k*[*P*] set proposed by Feldman [38, 76], where *P* denotes the bias (i.e., the number of input combinations for which the output is ‘True’) and is constrained to be odd, it was shown that NCFs yield the minimum average sensitivity (see Section 2.5) [37]. With that background, our goal here is to compare the Boolean complexity and average sensitivity of our three subtypes of EThF, namely NCF, IMR and BMR across varying number of external regulators. It is important to note that for a target node with *k* external regulators, a BMR rule corresponds to a (*k* + 1)-input EF, whereas an IMR rule corresponds to a *k*-input EF if *k* is odd and to a (*k* + 1)-input EF if *k* is even (see Section 3.1). Consequently, the range of bias values differs across various subtypes and that is why comparing the classes based on each bias individually is not suitable here. Instead, we compare the mean values of the two complexity measures – Boolean complexity and average sensitivity – computed over all unique BFs belonging to each subtype for different values of external regulators (*k*), going up to *k* = 9. We refer these two observables as the ‘averaged Boolean complexity’ and ‘averaged sensitivity’ respectively.

As can be readily verified from Eq. 3, the Boolean complexity of a *k*-input BF from the NCF class is always exactly *k*. In contrast, for a BMR regulated by *k* external regulators, the corresponding BF effectively has (*k* + 1) inputs. Therefore, the possible minimum Boolean complexity in this case is (*k* + 1). However, Fig. 3(a) reveals that the averaged Boolean complexity in the BMR class is substantially higher than this theoretical minimum for all *k >* 1. Furthermore, the deviation from the minimum increases with increasing *k*. To examine this in greater detail, we compute the Boolean complexity across all possible sign combinations of the *k* external regulators, for values of *k* up to 6 (see SI Fig. S4). It is evident from the figure that a BMR achieves the minimum possible Boolean complexity only under two specific conditions: when all regulators are either activatory or inhibitory. These two cases correspond precisely to the scenarios in which a BMR belongs to the class of NCFs (see SI text, Section 2, Corollary 2.2). For all other sign combinations, the Boolean complexity remains significantly higher than (*k* + 1). These observations suggest that, for any given *k*, the expected Boolean complexity of BMRs is significantly greater than that of NCFs. A similar pattern is observed for IMR. An IMR regulated by *k* external regulators corresponds to a *k*-or (*k* + 1)-input EF. As shown in Fig. 3(a), for any number of external regulators *k >* 1, the averaged Boolean complexity of IMRs consistently deviates from *k* and (*k* + 1), respectively. SI Fig. S4 further supports this observation by demonstrating that, for each *k* from 2 to 6, the Boolean complexity of IMRs never attains either of these minimal values. Interestingly, as *k* increases, the averaged Boolean complexities of IMRs and BMRs exhibit an alternating pattern (see Fig. 3(a)). Specifically, for odd values of *k*, BMRs tend to have higher averaged Boolean complexity than IMRs, whereas for even values of *k*, the reverse is observed.

**FIG. 3.**
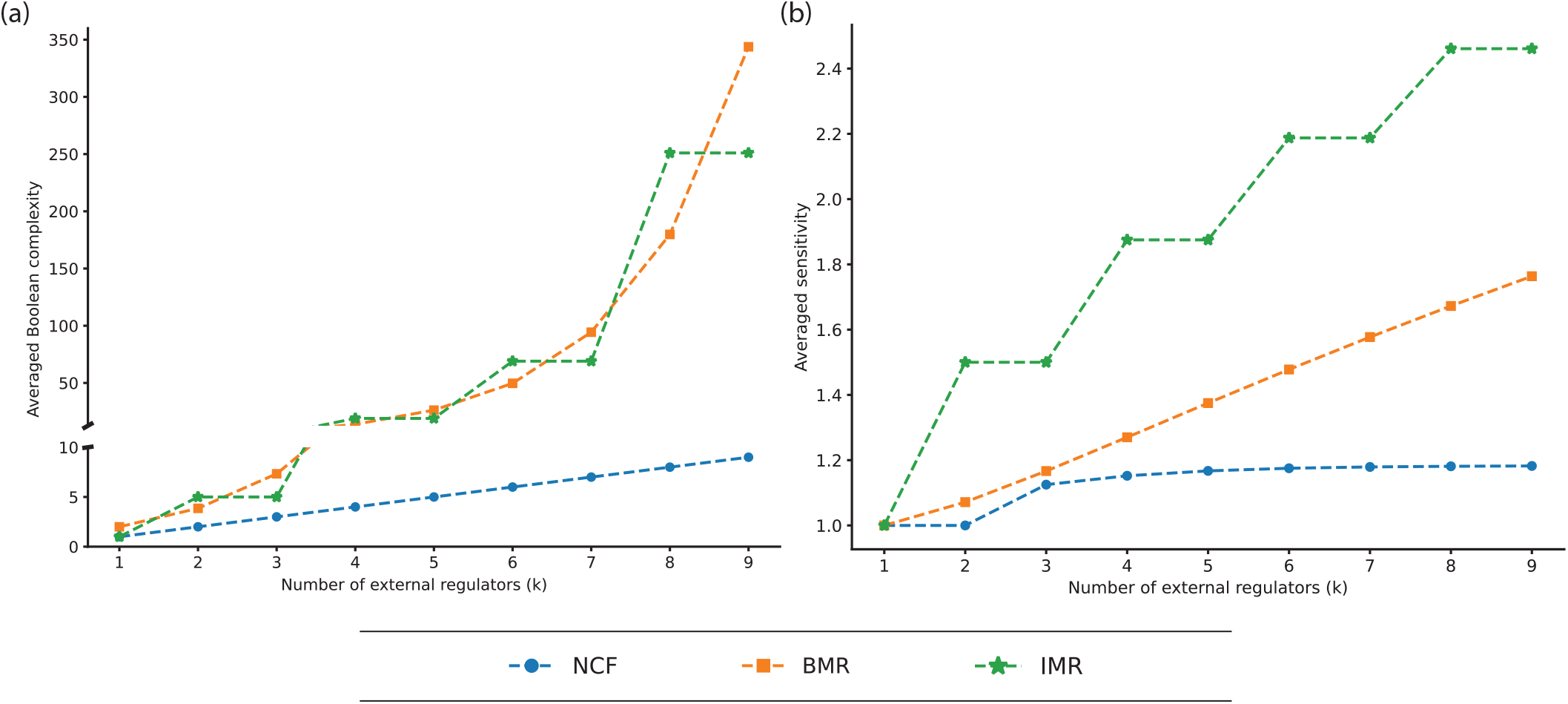
Values of averaged Boolean complexities and averaged sensitivities for NCFs, IMRs and BMRs. **(a)** The mean of the Boolean complexities of all unique NCFs, BMRs and IMRs across different number of external regulators *k* of a target node. A break has been introduced in the *y*-axis to accommodate the large gap between the values for NCFs and those for the TMRs. **(b)** The mean of the average sensitivities of all unique NCFs, BMRs and IMRs across different number of external regulators *k* of a target node.

We now turn to comparing averaged sensitivities of the three subtypes across different numbers of external regulators *k*. For *k >* 1, this measure reveals a consistent trend: NCF *<* BMR *<* IMR (see Fig. 3(b)). As *k* increases, the averaged sensitivity for NCFs grows slowly. In contrast, for TMRs the increase is much steeper. SI Fig. S5 shows the average sensitivity for all possible sign combinations of regulators for each subtype, for *k* up to 6. One interesting observation from this figure is that, for *k >* 1 and given signs for the regulators, there are multiple distinct average sensitivity values associated with the NCF class. This variation arises because, for a fixed sign pattern of the regulators, there exist NCFs with different bias values which may result in different values of the average sensitivity. In contrast, TMRs exhibit only a single average sensitivity value per sign combination. This is because there is a unique TMR for a given sign combination of the regulators, and thus, there is a unique bias value (see SI text, Properties 2.1, 2.2, and 2.8). Since average sensitivity is invariant under permutations of the inputs, the value remains fixed. It is also worth noting that, for several sign combinations, BMRs have lower average sensitivity than many NCFs corresponding to the same sign pattern of the regulators. However, in every case, except when all regulator signs are the same, there always exist some NCFs with lower average sensitivity than the corresponding BMR. On the other hand, IMRs consistently show the highest average sensitivity across all sign combinations. Importantly, this is not just a trend within these three subtypes; in fact, IMRs achieve the maximum average sensitivity among all *k*-input BFs in the UF family (see SI Text, Property 2.10).

These findings show that although TMRs offer a tempting simplicity for modeling GRNs, their underlying logic can be quite complex. On average, both IMRs and BMRs exhibit higher Boolean complexity and average sensitivity than NCFs – BFs that are highly enriched in the empirical datasets. Moreover, for any fixed sign combination of a node’s regulators, IMRs achieve the highest average sensitivity among all sign conforming UFs. Because sensitivity reflects robustness to input perturbations, this renders using IMRs the least robust choice. In sum, TMRs yield logic that is significantly more complex and sensitive to noise than NCFs.

### 3.4. Choosing threshold majority rules over nested canalyzing functions impairs recovery of the dynamical outcomes in biological Boolean networks

Now we address a question about the dynamical implications of using TMRs when modeling BNs: Can the TMRs substitute for the BFs of the published models while preserving key dynamical behaviors? From the 245 models in the BBM repository [55], our multi-step filtration pipeline yielded 24 high-confidence models that satisfy different structural and dynamical criteria (see Section 2.6). Each of these 24 selected published models recover the literature-reported biological attractors. We now ask, if every BF in these networks is replaced by TMRs respecting the activatory or inhibitory nature of the interactions, how well do the resulting alternative models recover the biological attractors and maintain their basin size distributions? To quantify this, we compute the quantities ARS_*alt*_, *D*_*model*_ and *D*_*alt*_ (see Section 2.7). For each of the 24 published models, we generate the two unique alternative variants that respect the signs of the interactions, one with IMRs and one with BMRs. Then, in each case, we simulate all three models to identify their complete attractor sets and associated basins of attraction.

#### Threshold majority rules perform poorly in recovering the biologically meaningful attractors

The attractors in biological Boolean networks mirror key biological outcomes such as cell-types, apoptosis, proliferation or cell-cycle, so a useful model must be able to embody them. Fig. 4 displays the attractor recovery score for the two alternative models when using IMRs and BMRs at each node, for each of the 24 network models. A score below 1 indicates inferior performance relative to the published model, with values near zero indicating an almost complete failure to recover the biologically meaningful attractors. Clearly, for a majority of the networks, both IMR- and BMR-based models yield scores near or equal to zero, demonstrating their inability to capture the biologically desired dynamical behavior. A score of exactly one means that the alternative model matches the performance of the published one. This occurs only for a couple of small networks when using IMR rules. For network 7 (network numbers are the IDs that uniquely identify each network of the BBM dataset), the extremely small state-space may allow even random rules to recover the biological attractors by chance. For the other network (network 110), that consists of nine nodes, six nodes have in-degree one. All the BFs assigned in the published model for this network are NCFs. Since, for *k* = 1, IMRs coincides exactly with NCFs, replacing the rules by IMRs leaves two thirds of the original logic intact. As the network size and average connectivity grow, the performance when using IMRs rapidly deteriorates. On the other hand, BMR performs worse than the published model across all cases, reinforcing the result that simple TMRs are generally insufficient to preserve the attractor structure of biological networks.

**FIG. 4.**
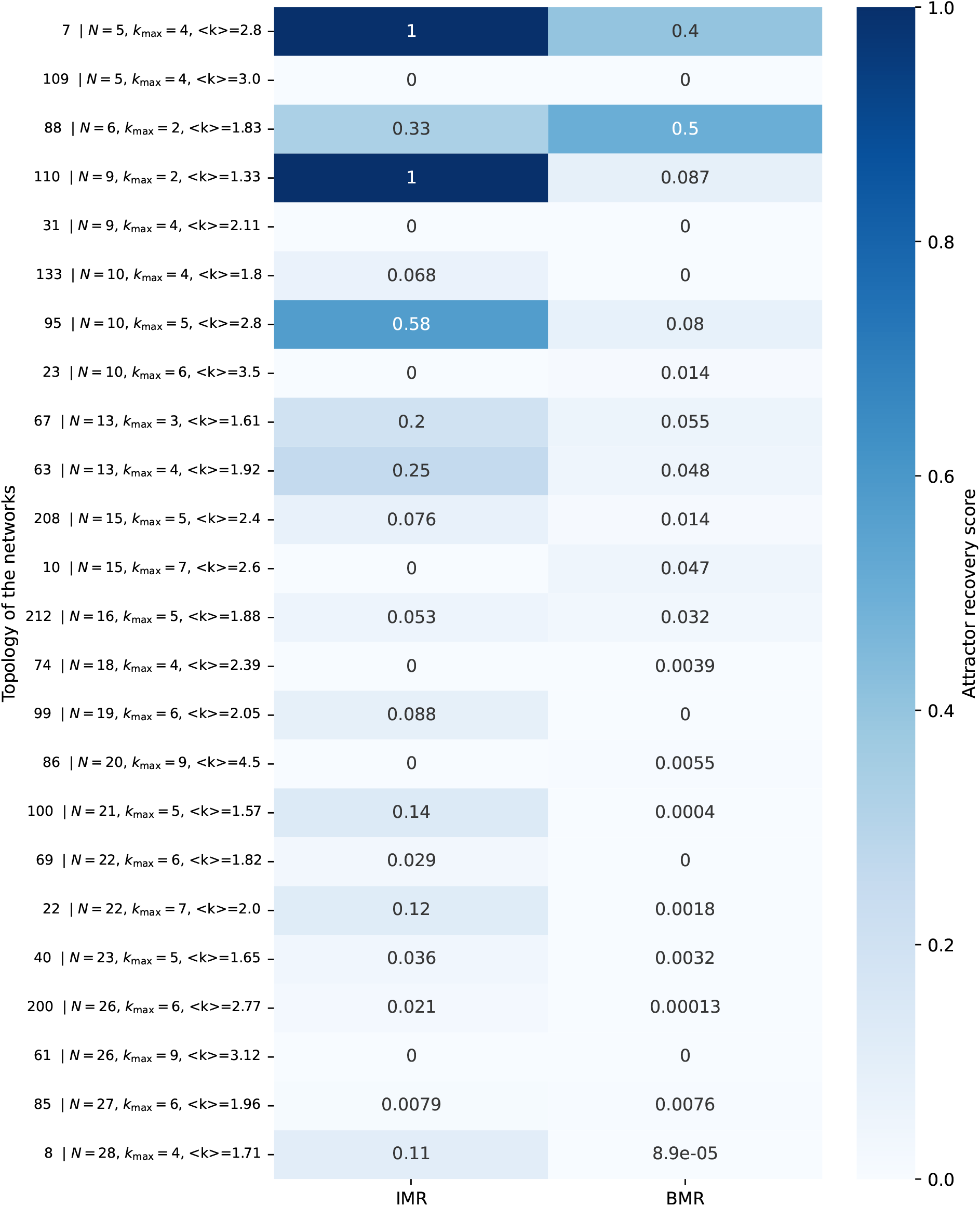
Attractor recovery scores when using IMRs and BMRs. Each row corresponds to a specific network from the BBM dataset. On the far left, the number indicates the network ID as assigned in the BBM dataset. The next columns list the number of nodes (*N*), the maximum in-degree (*k*_max_), and the average in-degree (*⟨ k⟩*) of the network. Rows are ordered in ascending order of *N*, followed by *k*_max_, and then *⟨ k⟩*. This heatmap displays the attractor recovery score for the alternative models constructed using IMR and BMR variants of the published model. A score close to 0 indicates poor performance in recovering the biological attractors, whereas a score of 1 indicates that the corresponding TMR-based model performs equally well as the published one in terms of attractor recovery.

#### The basin fractions of biologically meaningful attractors when using threshold majority rules are far from the desired target

Next we evaluate how closely the basin fraction distribution of the published models and their corresponding IMR and BMR variants align with a reconstructed ‘gold standard’ (see Section 2.7). This comparison is quantified using the Jensen–Shannon (JS) distance which measures the divergence between the basin size distribution of a model and the idealized distribution derived from the published model, considering only the biological attractors. The JS distance ranges from 0 to 1, where a value of 0 indicates perfect alignment with the gold standard distribution, and a value close to 1 indicates a very large deviation thereof. As can be seen in Fig. 5, several published models themselves exhibit considerable distances from the gold standard, largely due to the presence of spurious attractors whose basins occupy a substantial portion of the state space. This indicates that some published models are unsatisfactory, a point that we have emphasized in one of our previous works [77]. Nevertheless, models constructed using TMRs consistently show poorer alignment with the gold standard compared to the published models (i.e., *D*_*T MR*_ *> D*_*model*_), as indicated by their uniformly high JS distances across most cases. The sole exception is model 88 with IMR rules, which has a network size of only 6 nodes. Overall, these findings underscore a key limitation of using TMRs for modeling biological networks in general. Although they represent a very simple modeling choice, such simplicity generally comes at the cost of failing to preserve essential dynamical features of biological systems.

**FIG. 5.**
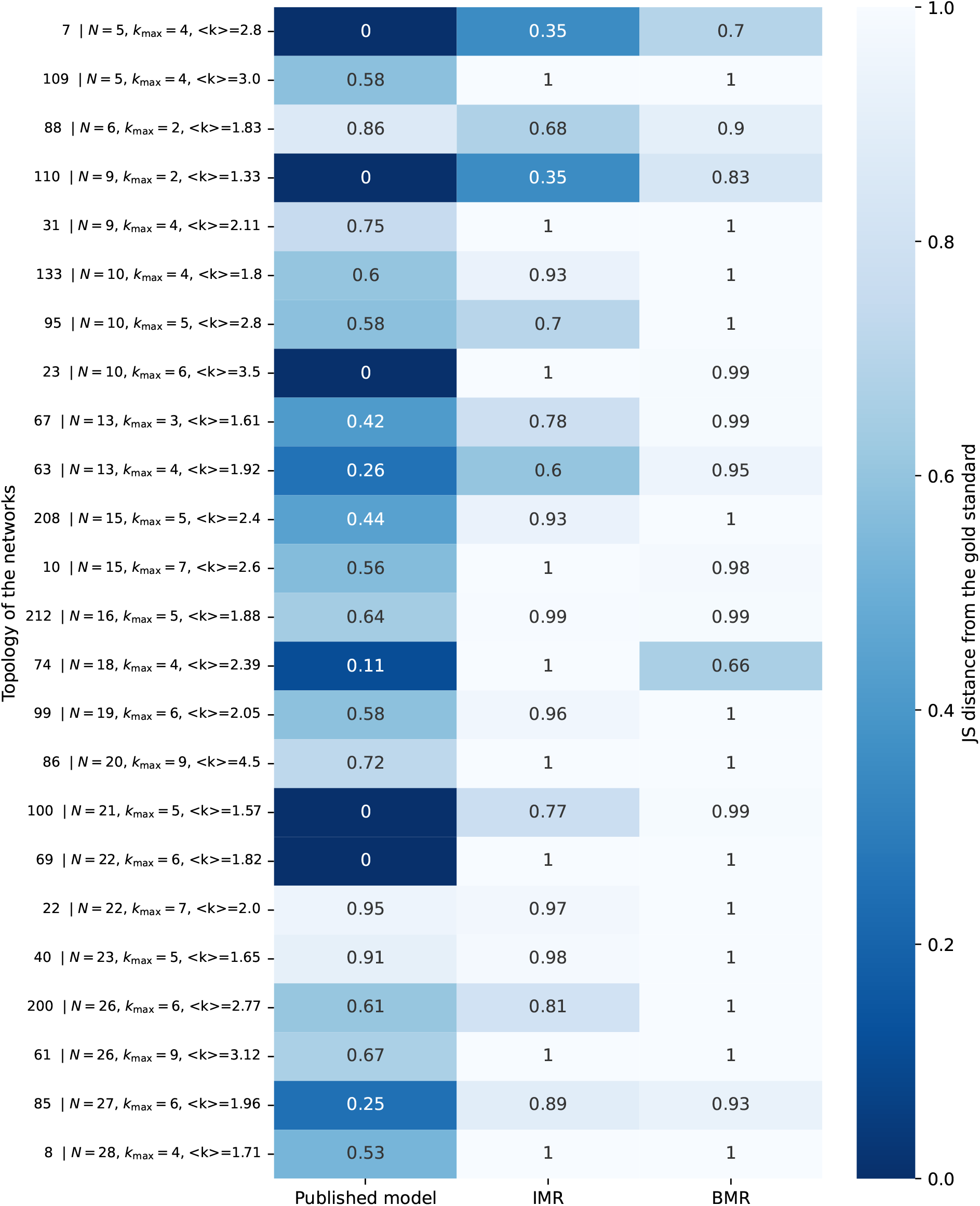
JS distance to gold standard when based on the fractions covered by basins of attraction. Each row corresponds to a specific network from the BBM dataset. On the far left, the number indicates the network ID as assigned in the BBM dataset. The next columns list the number of nodes (*N*), the maximum in-degree (*k*_max_), and the average in-degree (*⟨ k⟩*) of the network. Rows are ordered in ascending order of *N*, followed by *k*_max_, and then *⟨ k⟩*. This heatmap shows the Jensen-Shannon (JS) distance between the distribution of the published, IMR, and BMR variant models’ basin fractions and its corresponding gold standard. A JS distance of 0 indicates perfect overlap with the gold standard, while values closer to 1 reflect increasingly divergent distributions. Lower values therefore indicate better preservation of the original basin size structure.

#### Substituting non-nested canalyzing functions with suitable nested canalyzing functions in published models can yield superior dynamical outcomes

It has been shown that NCFs are highly enriched in empirical datasets of BFs [37, 54]. Accordingly, it is not surprising that in 20 of the 24 published models considered in this study, every assigned BF is NCF. Nevertheless, the networks with IDs 61, 69, 95 and 212 each include at least one non-NCF in their original published form. This observation leads us to two questions. First, can we replace the non-NCFs with appropriate sign conforming NCFs (scNCFs) while leaving existing NCFs unchanged and still recover all biological attractors? Second, if such replacements exist, could some of the resulting models achieve attractor recovery scores equal to or higher than those of the published models, and yield JS distances equal to or lower than those of the published models?

Because a given sign pattern can correspond to multiple scNCFs, we explored the possible replacements in each of the four networks while also satisfying the biological attractors [77, 78]. We found that the total number of candidate models satisfying these constraints is 686244 for network 61, 96 for network 69, 2116 for network 95 and 48 for network 212. For network 61, we randomly sampled 5000 of the 686244 models, then exhaustively simulated each of them, and finally computed the values of ARS_*repl*_ and *D*_*repl*_, where ARS_*repl*_ denotes the attractore recovery score of an NCF-replaced model and *D*_*repl*_ denotes the JS distance previously described (see Section 2.7). Remarkably, 1640 of the sampled models matched or exceeded the ARS_*model*_ and 4064 models matched or showed lower values than *D*_*model*_. For the remaining three networks we performed exhaustive simulations of all candidate models. In every case we identified a subset of those candidate models for which ARS_*repl*_ *≥* ARS_*model*_ and *D*_*repl*_ *≤ D*_*model*_. Indeed, nearly one hundred percent of the candidate models outperformed the TMR-based alternative models on these metrics. A complete breakdown of model counts for each network is provided in the SI Table S11.

Overall, given our comparisons of dynamical properties of biological Boolean networks when using TMRs rather than NCFs, all results point to serious failures when choosing TMRs. Note that unlike BFs drawn from the class of NCFs where a single sign configuration admits multiple candidate BFs and one can often select alternatives that preserve the known biological attractors, TMRs offer no such flexibility. Indeed, once the network topology is fixed, the TMR at each node is uniquely determined, producing a fixed set of attractors that may omit the biologically relevant ones. Even when TMRs recover the biological attractors, they often generate a large number of spurious attractors; furthermore, their basin of attraction fractions rarely align with the desired ones. Consequently, it seems generally inappropriate to choose TMRs when modeling biological regulatory networks.

### 3.5. Comparison of dynamical properties in random networks when using different subtypes of threshold functions

Next we analyzed how the choice between the three different ThF subtypes (scNCF, BMR and IMR) can affect the dynamical stability of *random* Boolean networks (RBNs). For that, we first consider 12-node RBNs with varying mean connectivity under the P-P topology (see Section 2.9). We find that the scNCF models consistently yield the fewest attractors, BMR and IMR models producing an order of magnitude more, with BMR exhibiting the highest average attractor count (see Fig. 6). This proliferation of attractors in BMR models has been noted previously [79, 80], and it was also found that there is an abundance of fixed point attractors for these models. Next, we consider the bushiness and convergence rate stability measures [71] of the STGs associated with each type of model, wherein high values correspond to stable dynamics. Models using IMRs exhibit the lowest average *G*-density (see Section 2.8), whereas models using scNCFs and BMRs both yield distributions with similarly high mean values. Notably, BMR shows a much wider range of *G*-density whereas the *G*-densities for all scNCF models are well clustered at higher levels. In contrast, the measure of convergence rate *λ*_*GoE*_ has its highest values for the BMR models, corroborating an earlier finding that BMR rules induce very short transients [80].

**FIG. 6.**
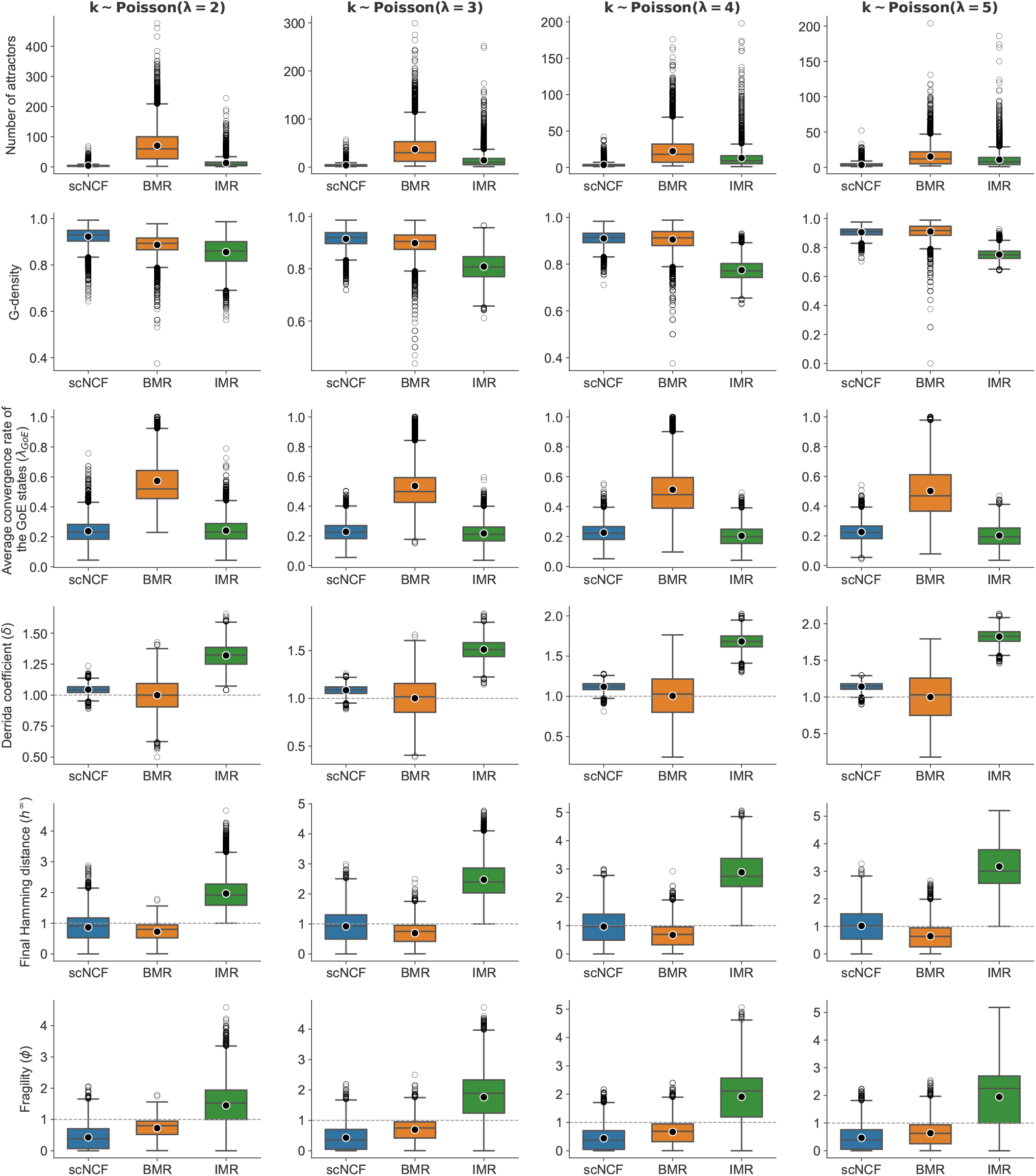
Distributions of stability measures for different subtypes of ThFs under the P-P network topology with *N* = 12. Each row corresponds to one observable and each of the four columns represent different values of the mean in-degree parameter (*λ* = 2, 3, 4, 5). In each subplot, the three boxplots present the distributions of the observables across three subtypes of ThFs: scNCFs, BMRs and IMRs. Outliers in each distribution are shown as grey circles while the black dots outlined in white denote the mean of each group.

Turning to damage spreading metrics, both *δ* and *h*^*∞*^ place the mean of the distributions for scNCF and BMR models very close to the critical boundary at 1 (*δ* = 1 and *h*^*∞*^ = 1, respectively), whereas the distributions for IMR models clearly fall in the chaotic regime. Notably, although the average *δ* for BMR is closer to the critical line than that of scNCFs, BMRs exhibit a much wider spread of *δ* values, reflecting a greater sensitivity to the underlying network topology. Finally, we consider the meaure *ϕ*, the phase-insensitive analogue of *h*^*∞*^. We find that scNCFs yield the lowest mean *ϕ*, BMRs are intermediate, and IMRs have the highest mean, well above the crtical threshold at *ϕ* = 1. The means for scNCF and BMR models both lie in the ordered regime. All these ordering patterns for each observable remain consistent as one varies the average connectivity (*λ*) over the values 2, 3, 4 and 5.

The qualitative patterns observed for 12-node networks persist when we analyze larger networks with 16-nodes (see SI Fig. S6). In this case, the differences in the distributions of number of attractors and *G*-density become even more noticeable for networks with higher connectivity. Across all scenarios, IMRs consistently perform the worst, while scNCFs and BMRs behave quite similar in most cases. These trends also hold when we switch from the P-P to the R-R topology (see SI Figures S7 and S8). One notable nuance appears in the distribution of the number of attractors. Under the P-P topology, BMRs produce the highest average attractor count for every tested value of the average connectivity. Under the R-R topology, where the in- and out-degree for each node is fixed (*k*_*in*_ = *k*_*out*_), the mean attractor count under BMR rules is maximum when *k*_*in*_ is odd. When *k*_*in*_ is even, IMR models take the lead in the average attractor count. In all cases, scNCF models maintain the fewest attractors. Lastly, the behaviour of all other stability measures under the R-R topology matches closely with that seen under P-P.

These results illustrate why we use multiple stability metrics: different measures can change the stability ranking of rule types. Nevertheless, by nearly all criteria, IMRs induce dynamical instability to the system while scNCFs and BMRs often tie and drive the dynamics more toward the stable regime. For some measures, such as number of attractors and *ϕ*, scNCFs clearly do best, and for *λ*_*GoE*_, BMRs stand out. Even though BMRs appear to drive the system toward stable dynamics, our primary concern in the context of modeling real biological networks remains the accurate recovery of known biological attractors (as discussed in Section 3.4). If a logic rule cannot achieve this fundamental requirement, its superior stability is of little practical utility. Finally, it is important to recognize that, even though we start with the same nominal network wiring, applying different types of rules can alter the effective architecture. Under scNCFs, the working topology matches the original exactly. However, with TMRs, a node that originally has *k* regulators may functionally behave as if it has *k −* 1, *k, k* + 1 or even 0 inputs, depending on the parity of *k*, the presence and signs of self-regulations, and the specific type of TMR used.

## 4. DISCUSSION AND CONCLUSIONS

Even for modestly sized GRNs with moderate to high connectivity, the number of Boolean models that reproduce a given set of biological attractors can be astronomically large, whether or not we confine our search within an exponentially smaller subspace of BFs such as NCFs to capture the combinatorial regulatory logic at single gene level. For instance, in a previous study [77], we showed that an 18-node GRN admits billions of such models, each of which recovers the biological attractors. This abundance makes it hard to decide which model or set of models is the ‘best’ for a specific biological system, particularly when the criteria for that choice may differ depending on the property of interest. Faced with this daunting model selection problem, modelers often turn toward TMRs. Even though TMRs are attractive for their uniformity and ease of implementation, one must ask whether this simplicity incurs acceptable compromises in biological realism. If the answer is no, the widespread use of such templates must be reconsidered. To address this, we have systematically analyzed TMRs and compared them against NCFs, a class of BFs that is highly enriched in the empirical datasets.

We summarize our findings in this context as follows. First, IMRs tend to produce BFs with even bias, and both TMR variants (IMRs and BMRs) allow very limited scope for canalyzation, standing in stark contrast to the prevalence of odd-biased BFs and the enrichment of NCFs in empirical datasets. Equally concerning is TMRs’ treatment of self-inhibitions. BMRs ignore self-inhibitions entirely and IMRs do so whenever the number of regulators is even; this is despite the critical role played by negative auto-regulation in biological GRNs. Second, our survey of three empirical datasets of BFs reveals that IMRs are underrepresented relative to their theoretically expected frequency. Although BMRs initially appeared to be enriched, this effect was largely from those BFs that are also NCFs; non-NCF BMRs are actually as scarce as IMRs. Third, we found that the subtype NCF yields lower averaged Boolean complexity and averaged sensitivity for any given number of regulators compared to either of the TMR types. Moreover, the IMRs attain the maximum possible average sensitivity among all UFs. Fourth, we asked whether TMRs still serve as a practical replacement for BFs in reconstructed Boolean GRN models. To explore this, by applying a rigorous filtration pipeline, we selected high-confidence reconstructed biological models whose whole state space can be exhaustively computed. By simulating each network with its BFs swapped for IMR or BMR subtypes, we found that both variants consistently showed poor performance in the recovery of biological attractors and their expected basin size distributions in almost all cases. In contrast, when we substituted any non-NCF BFs present in the published model with NCFs, we always identified some alternative models that matched or outperformed the published model on both biological attractor recovery and the basin fraction distributions. Finally, we evaluated network stability when using the different rule types via a suite of dynamical metrics. We observed in RBNs that typically both the BMRs and NCFs perform similarly and keep the dynamics close to the ordered or critical regime, whereas IMRs exhibit dynamics skewed toward the chaotic regime. Furthermore, both TMR types tend to produce a large number of attractors, many of which have very small basins. This phenomenon was also observed in well-known cell-cycle Boolean models of *S. cerevisiae* [31] *and S. pombe* [32], both of which used BMRs, where attractors with small basins were deemed biologically irrelevant and were discarded, which of course is not very satisfactory.

While we were working on this study, a preprint [81] appeared that also evaluated the two types of TMRs, which they referred to as the Ising (i.e., IMR) and 01 (i.e., BMR) formalisms, respectively. They report that in small regulatory networks with low connectivity, TMRs reproduce dynamics similar to real systems – an observation that aligns to some extent with our findings that TMRs typically yield higher ARS in small networks. Based on measures of function and state space similarity, for up to six regulators, they noted 65 *−* 80% mean similarity between TMRs and the biological rules. However, our analyses suggest that this overall similarity can be misleading. Although TMRs show similarity with the BFs in the empirical dataset based on their proposed measures, they often fail the critical task of recovering known biological attractors, which must remain the prime benchmark for any GRN model. Moreover, the preprint did not examine the intrinsic complexity or robustness of TMRs. We did this by measuring their Boolean complexity and average sensitivity, comparing to the case of NCFs, and found that TMRs are far more complex and sensitive. Using ensembles of RBNs, we also showed how IMRs drive network dynamics into a chaotic regime. The preprint [81] further proposes the modified versions, Ising* and 01*, claiming improved performance. We tested these variants across our 24 selected models and observed only marginal gains: cases with an ARS of 0 remained zero, and gains elsewhere were typically minimal (results not shown). Although Ising* functions produce odd bias when *k* is a power of 2, they still fail to be canalyzing for *k >* 2. On the other hand, 01* continues to render certain regulations ineffective. In light of these findings, neither the original nor the modified TMRs appear to offer a reliable default logic for modeling GRNs.

In summary, our results show that TMRs, however simple, fall short as a general framework for modeling regulatory networks as they arise in biological systems. While they may perform satisfactorily in certain cases, their rigid, one-size-fits-all logic offers no flexibility and often misaligns with biology. In contrast, NCFs, another subtype of ThFs, offer multiple sign conforming choices, exhibit minimal Boolean complexity and average sensitivity, and recover biological attractors with a much higher success rate and greater dynamical stability. Furthermore, their consistent enrichment in empirical datasets hints an evolutionary preference toward these rules. We therefore recommend moving beyond TMRs – in spite of their being particularly convenient – because their drawbacks are just too severe when seeking biological realism. Rather than sacrificing realism for simplicity, the challenge ahead lies in developing efficient methods to navigate the biologically meaningful classes of functions (which can faithfully capture the combinatorial regulation at single gene level) and select the most appropriate model for each system.

## ACKNOWLEDGMENTS

Olivier C. Martin and Areejit Samal would like to acknowledge funding from Indo-French Centre for the Promotion of Advanced Research (IFCPAR/CEFIPRA) via Collaborative Scientific Research Programme (CSRP) project number 7004-1. Areejit Samal acknowledges funding from the Department of Atomic Energy, Government of India via Apex project to The Institute of Mathematical Sciences (IMSc), Chennai, India. IPS2 benefits from the support of Saclay Plant Sciences-SPS (ANR-17-EUR-0007). Priyotosh Sil would like to acknowledge IFCPAR/CEFIPRA for support through the award of a Raman-Charpak fellowship.

## AUTHOR CONTRIBUTION

Designed the research: P.S., O.C.M., A.S.; Performed the research: P.S., O.C.M., A.S.; Performed the computations: P.S.; Wrote the paper: P.S., O.C.M., A.S.

## DATA AND CODE AVAILABILITY

All data and codes needed to reproduce the results reported in the main text and supplementary information (SI) of this manuscript are deposited in GitHub and are available at: https://github.com/asamallab/TMRvsBioLogic.

