## Supplementary material for "Simple threshold-based Boolean rules fall short in capturing biological regulatory network dynamics": SI

---

* To whom correspondence should be addressed:


### SUPPLEMENTARY TEXT

### 1. TYPES OF BOOLEAN FUNCTIONS

*Effective functions (EFs)*

Consider a Boolean function (BF) $f : \{0,1\}^k \to \{0,1\}$. The BF $f$ is *effective* [1] if every input bit $i \in \{1,2,\ldots,k\}$ can influence the output in at least one of the assignments of the remaining variables. Formally,

$$\forall\, i \in \{1,\ldots,k\},\ \exists\, \mathbf{x} \in \{0,1\}^k \text{ with } x_i = 0 \quad \text{s.t.} \quad f(\mathbf{x}) \neq f(\mathbf{x}+\mathbf{e}_i), \tag{1}$$

where $\mathbf{e}_i$ is the $i^{\text{th}}$ standard basis vector (all zeros except a '1' in position $i$). Effectiveness in all inputs is necessary for biological relevance but may not be sufficient.

*Unate functions (UFs)*

A BF $f$ with $k$ inputs is called *unate* [2] if each input variable $x_i$ is either monotone increasing (activatory) or monotone decreasing (inhibitory). Specifically, $x_i$ is monotone increasing if

$$\forall\, \mathbf{x} \in \{0,1\}^k \text{ with } x_i = 0, \quad f(\mathbf{x}) \ \leq\ f(\mathbf{x}+\mathbf{e}_i) \tag{2}$$

and monotone decreasing if

$$\forall\, \mathbf{x} \in \{0,1\}^k \text{ with } x_i = 0, \quad f(\mathbf{x}) \ \geq\ f(\mathbf{x}+\mathbf{e}_i) \tag{3}$$

*Canalyzing functions (CFs)*

A BF $f$ with $k$ inputs is said to be *canalyzing* [3] in input $i$ if there exist $a, b \in \{0,1\}$ such that

$$f(x_1, x_2, \ldots, x_{i-1}, a, x_{i+1}, \ldots, x_k) = b \tag{4}$$

for all choices of $x_j$ with $j \neq i$. Here '$a$' is the canalyzing input value and '$b$' the canalyzed output. The function $f$ is called a canalyzing function if it is canalyzing for at least one of its $k$ inputs.

*Read-once functions (RoFs)*

A $k$-input BF is called a *read-once* function (RoF) [4] if it can be written using only disjunction ($\vee$), conjunction ($\wedge$) and negation such that each of the variables appears exactly once. Equivalently, there exists a permutation $\pi$ of $\{1,2,\ldots,k\}$ and operator choice $\odot_i \in \{\wedge, \vee\}$ for each $i$ such that

$$f(\mathbf{x}) \ =\ X_{\pi(1)} \ \odot_1\ X_{\pi(2)}\ \odot_2\ \cdots\ \odot_{k-1}\ X_{\pi(k)}, \tag{5}$$

where each $X_{\pi(i)} \in \{x_{\pi(i)}, \overline{x}_{\pi(i)}\}$. For brevity, we omitted the parentheses that fully specify the evaluation order in the above equation.

### 2. PROPERTIES OF THE IMR AND BMR SETS

**Property 2.1.** For an IMR with $k$ regulators, where $k$ is odd, the BF equal to that IMR exhibits a bias of $2^{k-1}$ regardless of the sign combination of the regulators.

*Proof.* The bias of a BF is given by the number of input combinations that lead to 1 or 'on' as the output. Since, a BF with $k$ inputs has $2^k$ different input combinations, it would suffice to prove that the BF will have equal number of $+1$ ('on') and $-1$ ('off') in the output. Assume that we have $k$ inputs, with $k$ odd and each input $x_j \in \{-1, +1\}$. Further, each input is assigned with a sign $\sigma_j \in \{-1, +1\}$ depending on its activatory or inhibitory characteristics. Then, the output of the BF is given by:

$$f(x_1, x_2, \ldots, x_k) = \begin{cases} +1, & \text{if } S > 0 \\ -1, & \text{if } S < 0 \end{cases}$$

where, $S = \sum_{j=1}^{k} \sigma_j x_j$.

Let us define a mapping, $\phi : \{-1, +1\}^k \to \{-1, +1\}^k$, $\phi(x_1, x_2, \ldots, x_k) = (-x_1, -x_2, \ldots, -x_k)$. Clearly, the map is a bijection since each element of $\{-1, +1\}^k$ is uniquely mapped to an element in $\{-1, +1\}^k$ and $\phi^2(x) = x \; \forall x \in \{-1, +1\}^k$.

For an element $x = (x_1, x_2, \ldots, x_k) \in \{-1, +1\}^k$, $S(x) = \sum_{j=1}^{k} \sigma_j x_j$. Now, $S(\phi(x)) = \sum_{j=1}^{k} \sigma_j(-x_j) = -\sum_{j=1}^{k} \sigma_j x_j = -S(x)$. So, whenver $S(x) > 0, S(\phi(x)) < 0$ and vice versa. Since, $k$ is odd and each $x_i$ can be either +1 or -1, $S$ can never be zero. Thus, $\{-1, +1\}^k$ can be split into two non-overlapping sets:

$$S_+ = \{x \in \{-1, +1\}^k \mid S(x) > 0\} \quad \text{and} \quad S_- = \{x \in \{-1, +1\}^k \mid S(x) < 0\}.$$

Now, since $\phi$ is a bijection, it maps each element in $S_+$ to a unique element in $S_-$ and vice-versa. So, the sets $S_+$ and $S_-$ have same cardinality, and hence, the number of times $S > 0$ is $2^k/2 = 2^{k-1}$. Thus, the bias of the BF is $2^{k-1}$. □

**Property 2.2.** If an IMR rule is assigned to a node in a Boolean network with $k$ regulators, where $k$ is even, then the bias of the EF equal to that rule depends on the existence of a self-regulation at that node and its sign: (a) If the node is not self-regulating (i.e., it is not among its own inputs), then for any given sign combination, the IMR corresponds to a $(k+1)$-input EF with a bias of $2^k$. (b) If the node has a self-regulatory activatory edge, then the IMR corresponds to a $k$-input EF with a bias of $2^{k-1}$ for $k \geq 4$. (c) If the node has a self-regulatory inhibitory edge, then the IMR is equal to a a $(k-1)$-input EF with bias of $2^{k-2}$.

*Proof.* (a) If $k$ is even, then there exists $x = (x_1, x_2, \ldots x_k)$ such that $S$ (as defined in Property 2.1) equals to 0 and in that case it will depend on the its own state itself (say, $x_{k+1}$). Let us define the following sets:

$$A = \{x \in \{-1, 1\}^k \mid S(x) > 0\}$$
$$B = \{x \in \{-1, 1\}^k \mid S(x) < 0\}$$
$$C = \{x \in \{-1, 1\}^k \mid S(x) = 0\}$$

If $|A| = a, |B| = b, |C| = c$ then $a + b + c = 2^k$. Each element of $A$, $B$ and $C$ will contribute two, zero and one 1's in the output, respectively. So, the bias of the corresponding BF is $2a + c$. Now, for any $x$,

$$S(-x) = \sum_{j=1}^{k} \sigma_j(-x_j) = -S(x)$$

This means if $S(x) > 0$, then $S(-x) < 0$ and vice-versa. So, due to symmetry, $a = b$ (can be shown as in Property 2.1) and hence, $c = 2^k - 2a$. So, the bias of the equivalent BF is $2a + c = 2a + (2^k - 2a) = 2^k$.

(b) Now, suppose there exists a self-activatory edge. Without loss of generality, let us assume that $x_k$ represents the node with the self-regulation with an activatory effect. In this case, when the signed sum $S = 0$, the output will depend on the value $x_k$. Since the self-regulation is activatory, the signed sum can be written as $S(x) = \sum_{j=1}^{k-1} \sigma_j x_j + \sigma_k x_k = T + x_k$. Since $T$ is sum of an odd number of elements, namely $(k-1)$, where each element is either +1 or -1, the possible values that $T$ can attain are: $-(k-1), -(k-3), \ldots, -1, 1, \ldots, (k-3), (k-1)$. Now, consider two input configurations: $a = (a_1, a_2, \ldots a_{k-1}, 1)$ and $a' = (a_1, a_2, \ldots a_{k-1}, -1)$ such that the sum $T = \sum_{j=1}^{k-1} \sigma_j a_j = 1$. For the configuration $a$, the total signed sum becomes

$$S(a) = T + x_k = 1 + 1 = 2 > 0$$

Hence, the output is 1. Again, for the configuration $a'$, we have

$$S(a') = T + x_k = 1 - 1 = 0$$

Since, the signed sum is 0, the output is determined by $x_k$, leading to an output $-1$. This demonstrates that the self-regulation is effective. Therefore, it is possible to map a $k$-input (with even $k$) IMR with activatory self-regulation to a $k$-input effective BF. This result holds for all even $k \geq 4$.

However, for $k = 2$, the presence of an activatory self-regulation leads to a $k$-input ineffective BF. Specifically, in this case the other input does not influence the output, i.e., if $x = (x_1, x_2)$ and $x_2$ represents the node with activatory self-regulation, then for both the sign combinations '++' and '-+', the output depends solely on $x_2$. The BF reduces to the EF $f(x_1, x_2) = x_2$, in both cases resulting in a bias equal to 1. Now, for any even $k \geq 4$, we aim to show that the bias of a BF equal to an IMR with an activatory self-regulation is exactly $2^{k-1}$.

Suppose $k \geq 4$ and we consider $S(x) = T + x_k$. Now, $T$ can take the following values: $-(k-1), -(k-3), \ldots, -1, 1, \ldots, (k-3), (k-1)$. Suppose $N_t$ denotes the count of $x$ for which $T$ can be equal to $t$. Then, by symmetry $N_t = N_{-t}$ and hence

$$\sum_{t=-(k-1)}^{-1} N_t = \sum_{t=1}^{(k-1)} N_t = 2^{k-1} \text{ since } \sum_{t=-(k-1)}^{(k-1)} N_t = 2^k$$

For $T \leq -3$, $S < 0$ for both values of $x_k$. So, for all $x$ satisfying $T \leq -3$, the output is $-1$. Again, for $T \geq 3$, $S > 0$ for both values of $x_k$. So, for all $x$ satisfying $T \geq 3$, the output is 1. Now, when $T = 1$, if $x_k = 1$, then the output becomes 1, but if $x_k = -1$, then the output is $-1$. Which means when $T = 1$, half of the times we get 1 as output. Again, when $T = -1$, if $x_k = 1$, the output becomes 1 as $S = 0$ and $x_k = 1$. If $x_k = -1$ then the output is $-1$. So, again we see that when $T = -1$, half of the times we get 1 as output. Thus, the bias of the equivalent BF would be equal to

$$\sum_{t=3}^{(k-1)} N_t + \frac{1}{2}(N_1 + N_{-1}) = \sum_{t=1}^{(k-1)} N_t = 2^{k-1} \text{ since } N_1 = N_{-1}.$$

(c) Now consider the case when the self-regulation is inhibitory. Let the self-regulation be on the node $x_k$; then the total signed sum becomes $S(x) = T - x_k$ where $T = \sum_{j=1}^{(k-1)} \sigma_j x_j$. As discussed in (b), for even $k$, $T$ can take the odd integer values in the range $[-(k-1), (k-1)]$. Now, we analyze the output behaviour for different values of $T$.

- $T \leq -3$ : For both $x_k = \pm 1$, we have $S < 0$, so the output is $-1$.
- $T \geq 3$: For both $x_k = \pm 1$, we have $S > 0$, so the output is 1.
- $T = 1$:
  - If $x_k = 1$, then $S = 0$, so the output is 1 (since $x_k = 1$).
  - If $x_k = -1$, then $S = 2$, so the output is also 1.

  Thus, the output is 1 regardless of the value of $x_k$.
- $T = -1$:
  - If $x_k = 1$, then $S = -2$, so the output is $-1$.
  - If $x_k = -1$, then $S = 0$, so the output is $-1$ (since $x_k = -1$).

  Thus, the output is $-1$ regardless of the value of $x_k$.

This analysis shows that for $T = \pm 1$, the output is independent of $x_k$, implying that the inhibitory self-regulation becomes ineffective. Thus, it can be shown that the bias will be $2^{k-1}$ using the same approach as in (b). However, since it can be reduced to a $(k-1)$-input EF by removing the ineffective self-regulation, the bias will be $\frac{2^{k-1}}{2} = 2^{k-2}$. □

**Property 2.3.** A BF equal to an IMR with $k$ regulators can never be canalyzing for any $k > 2$, regardless of the presence or absence of a self-regulation.

*Proof.* The IMR rule can be written as:

$$f(x_1,\ldots,x_k,x_t)=\begin{cases}+1, & \sum_{i=1}^{k}\sigma_i\,x_i\;>\;0,\\ x_t, & \sum_{i=1}^{k}\sigma_i\,x_i\;=\;0,\\ -1, & \sum_{i=1}^{k}\sigma_i\,x_i\;<\;0,\end{cases}$$

where $x_i \in \{-1,+1\}$, and $x_t$ denotes an additional variable. When $k$ is even and there is no self-regulation, $x_t$ acts as an external variable (representing an implicit self-regulation). If $k$ is even and one of the $k$ regulators is already a self-regulator, $x_t$ is considered as an internal variable which is the self-regulator. For odd $k$, $x_t$ can be treated either as an internal variable or as an external ineffective variable, since the signed sum is never zero when $k$ is odd.

We will show that for every $1 \leq r \leq t$ and every fixed value $a \in \{+1,-1\}$, there always exists two assignements of the other variables under which $f$ takes both values $+1$ and $-1$. We denote $\alpha_i = sgn(\sigma_i)$ for all $i$ then $\sigma_i \cdot \alpha_i = 1; \sigma_i \cdot (-\alpha_i) = -1$ for all $i$.

**Case 1**: Suppose $x_t$ is an internal variable. Suppose $x_r$ is a canalyzing variable with $x_r = a$

**Input 1:** $x_r = a, x_i = \alpha_i\ \forall\ i \neq r$. Then, the signed sum

$$S=\sum_{i=1}^{k}\sigma_i x_i=\sigma_r a+\sum_{i\neq r}\sigma_i x_i=\sigma_r a+\sum_{i\neq r}\sigma_i\alpha_i=\sigma_r a+(k-1)\geq -1+(k-1)>0 \text{ since } k>2$$

So, the output is $+1$ in this case.

**Input 2:** $x_r = a, x_i = -\alpha_i\ \forall\ i \neq r$. Then, the signed sum

$$S=\sum_{i=1}^{k}\sigma_i x_i=\sigma_r a+\sum_{i\neq r}\sigma_i x_i=\sigma_r a+\sum_{i\neq r}\sigma_i(-\alpha_i)=\sigma_r a-(k-1)\leq 1-(k-1)<0 \text{ since } k>2$$

So, the output is $-1$ in this case. So, none of the variables is a canalyzing variable.

**Case 2**: Suppose $x_t$ is an external variable and $x_r$ is a canalyzing variable with $x_r = a$. If $r \neq t$ then we can choose the same pair of inputs as in Case 1 to show that $x_r$ cannot be canalyzing. If $r = t$ then we choose the following two inputs.

**Input 1:** $x_{r=t} = a, x_i = \alpha_i\ \forall\ i \neq t$. Then, the signed sum

$$S=\sum_{i=1}^{k}\sigma_i x_i=\sum_{i=1}^{k}\sigma_i\alpha_i=\sum_{i=1}^{k}(+1)=k>0$$

So, the output is $+1$ in this case.

**Input 2:** $x_{r=t} = a, x_i = -\alpha_i\ \forall\ i \neq t$. Then, the signed sum

$$S=\sum_{i=1}^{k}\sigma_i x_i=\sum_{i=1}^{k}\sigma_i(-\alpha_i)=\sum_{i=1}^{k}(-1)=-k<0$$

So, the output is $-1$ in this case. So, none of the variables is a canalyzing variable.

This completes the proof that a BF which is equal to an IMR with $k$ regulators can never be a canalyzing function if $k > 2$ irrespective of whether there exists a self-regulation or not. □

**Property 2.4.** Consider a BF $f$ that corresponds to a BMR with $k$-regulators, one of which is a self-regulation. Then: (a) for $k > 2$, if the self-regulation is activatory, $f$ is effective; (b) for $k > 1$, if the self-regulation is inhibitory but not all other regulators are inhibitory, $f$ is ineffective, with the self-input being the ineffective input; (c) for $k > 0$, if all $k$ regulators are inhibitory, $f$ reduces to the constant 0 function; and (d) for $k = 1$ or 2, the function $f$ is effective only when all regulators are activatory.

*Proof.* (a) Consider a BMR with $k$ regulators $x_1, x_2, \dots x_k$. Without loss of generality let us assume $x_k$ is the target node, and hence, is the self-regulatory input. If we denote the signed sum by $S = \sum_{i=1}^{k} \sigma_i x_i$, then the function $f$ is defined by

$$f(x_1, x_2, \dots, x_k) = \begin{cases} 1, & \text{if } S > 0 \\ 0, & \text{if } S < 0 \\ x_k, & \text{if } S = 0 \end{cases}$$

First consider $x_i = 0$ for all $i < k$. Then toggling $x_k$ from 0 to 1 changes the output from 0 to 1. So, the input $x_k$ is effective. Now we want to show that all $x_j$ (for $1 \le j < k$) are effective. Since $k > 2$, $\exists\ m < k$ such that $m \notin \{j, k\}$. If all $x_i = 0\ \forall\ i \ne m, j, k$ then the signed sum becomes $S = \sigma_j x_j + \sigma_m x_m + x_k$. So, the pair $(\sigma_j, \sigma_m)$ can be equal to any tuple from the set $\{(1,1), (1,-1), (-1,1), (-1,-1)\}$. If $(\sigma_j, \sigma_m)$ is equal to $(1,1)$ or $(1,-1)$ then fixing $x_m = x_k = 0$ and toggling $x_j$ from 0 to 1 changes the output from 0 to 1. If $(\sigma_j, \sigma_m)$ is equal to $(-1,1)$ then fixing $x_m = 1, x_k = 0$ and toggling $x_j$ from 0 to 1 changes the output from 1 to 0. If $(\sigma_j, \sigma_m)$ is equal to $(-1,-1)$ then fixing $x_m = 0, x_k = 1$ and toggling $x_j$ from 0 to 1 changes the output from 1 to 0. This proves that $f$ is effective when the self-regulation is activatory.

(b) Suppose the self-regulator is inhibitory then $\sigma_k = -1$. Let us denote $T = \sum_{i=1}^{k-1} \sigma_i x_i$. Then, $S = T - x_k$.

If $x_k = 0$,

$$f(x_1, x_2, \dots, 0) = \begin{cases} 1, & \text{if } T > 0 \\ 0, & \text{if } T \le 0 \end{cases}$$

If $x_k = 1$,

$$f(x_1, x_2, \dots, 1) = \begin{cases} 1, & \text{if } T - 1 > 0 \\ 1, & \text{if } T - 1 = 0 \\ 0, & \text{if } T - 1 < 0 \end{cases}$$

$$= \begin{cases} 1, & \text{if } T > 0 \\ 0, & \text{if } T \le 0 \end{cases}$$

So, the output depends only on the signed sum of first $(k-1)$ variables, $T$. Thus, the self-regulation becomes ineffective.

Note that using the same approach as in $(a)$, it can be shown that for all $1 \le j < k$ $(k > 2)$ whenever $(\sigma_j, \sigma_k) \in \{(1,1), (1,-1), (-1,1)\}$, with a suitable chosen value of $x_m$, and $x_i = 0\ \forall i \ne k, m, j$, toggling the value of $x_j$ from 0 to 1 will change the output. So, in all such cases $x_j$ will be an effective input for all $1 \le j < k$. Now, if for all $j, m < k$, $(\sigma_j, \sigma_k) = (-1,-1)$, then we can prove that none of the inputs is effective and $f$ is the constant zero function. See (c) below for the explanation.

(c) If all the regulators are inhibitory, $\sigma_i = -1\ \forall\ 1 \le i \le k$. Then, the signed sum becomes,

$$S = -\sum_{i=1}^{k} x_i$$

If $x_i = 0\ \forall\ 1 \le i \le k$, then $S = 0$. So, the output is 0 because $x_k$ is 0. Now, if there exists at least one $j$ $(1 \le j \le k)$ such that $x_j = 1$, then $S < 0$. So, the output is 0. Therefore, $f$ is the constant 0 function and no input affects the output.

(d) Consider a BMR with one input (which here is obviously the self-input). If the self-regulation is inhibitory then from (c) it follows that $f$ is ineffective. If the self-regulation is activatory then $f$ is the identity function, i.e., $f(x_1) = x_1$ and hence effective. Now, consider a BMR with two inputs $x_1, x_2$ and for which $x_2$ is the self-regulator. Now, if $\sigma_2 = -1$ then using (b), $f$ will be ineffective. If $(\sigma_1, \sigma_2) = (-1, +1)$, $f(x_1, x_2) = x_2$, so $f$ is ineffective. If $(\sigma_1, \sigma_2) = (+1, +1)$, $f(x_1, x_2) = x_1 \vee x_2$. This is the only case for which $f$ is effective. Thus, for $k = 1, 2$, $f$ will be effective only when all the regulators are activatory. $\square$

**Property 2.5.** The bias of a BMR having $k$ inputs, all external, is given by the following expression

$$P^k_{BMR} = 2 \cdot \sum_{m=0}^{a-1} \binom{k}{m} + \binom{k}{a}$$

where $a$ is the number of activators among those $k$ regulators.

*Proof.* A BMR with $k$ external regulators can be mapped to a $(k+1)$-input BF. $a$ is the number of activators. Then there are $(k-a)$ inhibitors. Suppose $i$ and $j$ denote the number of active activators and active inhibitors, respectively. Then, the conditions $i > j, i = j$ and $i < j$, will contribute two, one and zero 1's, respectively in the list of outputs. Thus, the bias of the BMR is given by

$$P^k_{BMR} = \sum_{i=0}^{a} \sum_{j=0}^{k-a} \binom{a}{i} \binom{k-a}{j} \cdot w(i,j) \tag{6}$$

where,

$$w(i,j) = \begin{cases} 2 & \text{if } \ i > j \\ 1 & \text{if } \ i = j \\ 0 & \text{if } \ i < j \end{cases}$$

So, we need to evaluate the double sum when $i > j$ and when $i = j$.

**Case 1 ($i > j$):** We need to determine the value of

$$P_{pos} = \sum_{\substack{i>j \\ 0 \le i \le a \\ 0 \le j \le k-a}} \binom{a}{i} \binom{k-a}{j}$$

Consider

$$G(x) = \sum_{i=0}^{a} \sum_{j=0}^{k-a} \binom{a}{i} \binom{k-a}{j} x^{i-j} \tag{7}$$

Then $G(x)$ can be written as

$$\begin{aligned} G(x) = \left( \sum_{i=0}^{a} \binom{a}{i} x^i \right) \cdot \left( \sum_{j=0}^{k-a} \binom{k-a}{j} x^{-j} \right) &= (1+x)^a \cdot (1+x^{-1})^{k-a} \\ &= x^{-(k-a)} (1+x)^k \\ &= x^{(a-k)} \sum_{s=0}^{k} \binom{k}{s} x^s \\ &= \sum_{s=0}^{k} \binom{k}{s} x^{s+a-k} \\ &= \sum_{t=-(k-a)}^{a} \binom{k}{t+k-a} x^t \end{aligned}$$

From Eq. 7, the coefficient of $x^t$ in $G(x)$ is

$$C_t = \sum_{\substack{i-j=t \\ 0 \le i \le a \\ 0 \le j \le k-a}} \binom{a}{i} \binom{k-a}{j}$$

Since $i > j$ iff $t \geq 1$, by equating the sum of coefficients of the positive powers of $x$, we have

$$\sum_{\substack{i>j \\ 0\leq i\leq a \\ 0\leq j\leq k-a}} \binom{a}{i}\binom{k-a}{j} = \sum_{t=1}^{a} \binom{k}{t+k-a}$$

$$= \sum_{r=k-a+1}^{k} \binom{k}{r}$$

Finally, using the binomial symmetry $\binom{k}{r} = \binom{k}{k-r}$, we have

$$P_{pos} = \sum_{\substack{i>j \\ 0\leq i\leq a \\ 0\leq j\leq k-a}} \binom{a}{i}\binom{k-a}{j} = \sum_{m=0}^{a-1} \binom{k}{m} \tag{8}$$

**Case 2 ($i = j$):** Now, we need to determine the value of

$$P_{null} = \sum_{\substack{i=j \\ 0\leq i\leq a \\ 0\leq j\leq k-a}} \binom{a}{i}\binom{k-a}{j}$$

Suppose $i = j = d$. Then

$$P_{null} = \sum_{d=0}^{min(a,\ k-a)} \binom{a}{d}\binom{k-a}{d}$$

If $min(a, k-a) = a$,

$$P_{null} = \sum_{d=0}^{a} \binom{a}{d}\binom{k-a}{d} = \sum_{d=0}^{a} \binom{a}{a-d}\binom{k-a}{d}$$

Using Vandermonde's convolution, we get,

$$P_{null} = \binom{a+(k-a)}{a} = \binom{k}{a}$$

Similarly, if $min(a, k-a) = k-a$,

$$P_{null} = \sum_{d=0}^{k-a} \binom{a}{d}\binom{k-a}{d} = \sum_{d=0}^{a} \binom{a}{d}\binom{k-a}{(k-a)-d}$$

Again, using Vandermonde's convolution,

$$P_{null} = \binom{a+(k-a)}{k-a} = \binom{k}{k-a} = \binom{k}{a}$$

So, irrespective of $min(a, k-a) = k-a$ or $a$, the value of $P_{null}$ is given by

$$P_{null} = \binom{k}{a} \tag{9}$$

Using Equations 6, 8 and 9, we have

$$P^k_{BMR} = 2 \cdot P_{pos} + P_{null} = 2 \cdot \sum_{m=0}^{a-1} \binom{k}{m} + \binom{k}{a}$$

□

**Corollary 2.1.** Suppose the BF $f$ corresponds to a BMR with $k$ external regulators, out of which $a$ are activatory. Then, (a) $f$ is an NCF if and only if all the regulators are inhibitory or all are activatory; (b) $f$ is of odd bias if and only if $a \wedge \overline{k} = 0$, where $a \wedge \overline{k}$ indicates the bitwise AND operation of $a$ and $\overline{k}$; (c) All such $f$ will be of odd bias if and only if $k$ is a Mersenne number.

*Proof.* (a) If all the regulators are inhibitory then $a = 0$. So, using Property 2.5

$$P^k_{BMR} = \binom{k}{0} = 1$$

Since, the bias is 1, it indicates an AND function, and hence, an NCF.
Now, if all the regulators are activatory, then $a = k$. Using Property 2.5

$$P^k_{BMR} = 2 \cdot \sum_{m=0}^{k-1} \binom{k}{m} + \binom{k}{k}$$

Now, using the identity $\sum_{m=0}^{k} \binom{k}{m} = 2^k$ we have,

$$\begin{aligned} P^k_{BMR} &= 2 \cdot \left[2^k - \binom{k}{k}\right] + \binom{k}{k} \\ &= 2^{k+1} - 1 \end{aligned}$$

Since, the bias is $2^{k+1} - 1$, all except one of the output values are equal to 1, which indicates it to be an OR function and hence an NCF.
Lastly, in the mixed case, $f$ cannot even be canalyzing (see Property 2.7) and consequently it is not an NCF.

(b) From Property 2.5, it follows that the bias can be odd iff $\binom{k}{a}$ is odd. As a consequence of Lucas's theorem [5] we know that $\binom{k}{a}$ is odd iff every binary bit of $a$ is less than or equal to the corresponding bit of $k$ [6]. In other words, $\binom{k}{a}$ is odd iff $a \wedge \overline{k} = 0$, where $a \wedge \overline{k}$ indicates the bitwise AND operation of $a$ and $\overline{k}$ (i.e., writing $a$ and $k$ as binary strings of length $k$, one performs the bitwise AND operation between $a$ and $\overline{k}$, and then, one converts the output to integer and checks if is zero) [6].

(c) For a given $k$, all the BFs will be of odd bias iff for all $0 \leq a \leq k$, $\binom{k}{a}$ is odd. Now, from (b) it follows that it is possible only when all the binary bits of $k$ are 1, and that is possible only if $k$ is of the form $2^r - 1$ for some positive integer $r$, i.e., only when $k$ is a Mersenne number. □

**Property 2.6.** The bias of a $k$-input BF $f$ corresponding to a BMR with $k$ regulators – one of which is a self-regulation – is given by the following expression:

$$P^k_{BMR} = \sum_{m=0}^{a-1} \binom{k}{m} + \binom{k-1}{a-\delta}$$

where $a$ is the number of activators among those $k$ regulators, $\delta = 0$ if the self-regulation is activatory and $\delta = 1$ if the self-regulation is inhibitory.

*Proof.* Suppose the $k$ regulators of a node $x_r$ are $x_1, \ldots, x_{r-1}, x_r, x_{r+1}, \ldots, x_k$. The BMR at node $x_r$ can thus be represented as a BF with $k$ inputs. The output of this function is 1 under the following two conditions: (1) the signed sum of the inputs is positive; (2) the signed sum is 0 and $x_r = 1$. Each instance in which either of these conditions holds contributes a 1 to the output of the function. Therefore,

$$P^k_{BMR} = P_{pos} + P_{null}(x_r = 1)$$

The value of $P_{pos}$ remains identical to that derived in Property 2.5:

$$P_{pos} = \sum_{m=0}^{a-1} \binom{k}{m} \tag{10}$$

Now we need to evaluate the value of $P_{null}(x_r = 1)$, i.e., the number of input combinations for which the signed sum is 0 and $x_r = 1$. Suppose, $a, i, j$ denote the number of activators, number of active activators and number of active inhibitors, respectively. The signed sum is 0 when $i = j$, leading to

$$P_{null} = \sum_{j=0}^{min(a,k-a)} \binom{a}{j}\binom{k-a}{j}$$

Among these we need to count only the input combinations where $x_r = 1$. Since the self-regulation at $x_r$ can be activatory or inhibitory in nature, we consider these two cases separately.

**Case 1 ($x_r$ is activatory)**: Since, $x_r = 1$ and it is self-activatory, we need to choose $(j-1)$ other active activators from the remaining $(a-1)$ activator slots. So, in this case,

$$P_{null}(x_r = 1) = \sum_{j=0}^{min(a,k-a)} \binom{a-1}{j-1}\binom{k-a}{j} \tag{11}$$

If $min(a, k-a) = a$,

$$\begin{aligned} P_{null}(x_r = 1) &= \sum_{j=0}^{a} \binom{a-1}{j-1}\binom{k-a}{j} \\ &= \sum_{j=0}^{a} \binom{a-1}{a-j}\binom{k-a}{j} \\ &= \binom{k-1}{a} \text{ (using Vandermonde's convolution)} \end{aligned}$$

If $min(a, k-a) = k-a$,

$$\begin{aligned} P_{null}(x_r = 1) &= \sum_{j=0}^{k-a} \binom{a-1}{j-1}\binom{k-a}{j} \\ &= \sum_{i=-1}^{k-a-1} \binom{a-1}{i}\binom{k-a}{i+1} \\ &= \sum_{i=0}^{k-a-1} \binom{a-1}{i}\binom{k-a}{(k-a-1)-i} \\ &= \binom{k-1}{k-a-1} \text{ (using Vandermonde's convolution)} \\ &= \binom{k-1}{a} \end{aligned}$$

Thus, when $x_r$ is activatory,

$$P_{null}(x_r = 1) = \binom{k-1}{a}$$

**Case 2 ($x_r$ is inhibitory)**: Since, $x_r = 1$ and it is self-inhibitory, we need to choose $(j-1)$ other active inhibitors from the remaining $(k-a-1)$ inhibitor slots. So, in this case,

$$P_{null}(x_r = 1) = \sum_{j=0}^{min(a,k-a)} \binom{a}{j}\binom{k-a-1}{j-1} \tag{12}$$

If $min(a, k-a) = a$,

$$\begin{aligned} P_{null}(x_r = 1) &= \sum_{j=0}^{a} \binom{a}{j}\binom{k-a-1}{j-1} \\ &= \sum_{i=-1}^{a-1} \binom{a}{i+1}\binom{k-a-1}{i} \\ &= \sum_{i=0}^{a-1} \binom{a}{(a-1)-i}\binom{k-a-1}{i} \\ &= \binom{k-1}{a-1} \text{ (using Vandermonde's convolution)} \end{aligned}$$

If $min(a, k-a) = k-a$,

$$\begin{aligned} P_{null}(x_r = 1) &= \sum_{j=0}^{k-a} \binom{a}{j}\binom{k-a-1}{j-1} \\ &= \sum_{j=0}^{k-a} \binom{a}{j}\binom{k-a-1}{(k-a)-j} \\ &= \binom{k-1}{k-a} \text{ (using Vandermonde's convolution)} \\ &= \binom{k-1}{a-1} \end{aligned}$$

Thus, when $x_r$ is inhibitory,

$$P_{null}(x_r = 1) = \binom{k-1}{a-1}$$

A single formula can then summarize the two cases:

$$P^k_{BMR} = \sum_{m=0}^{a-1} \binom{k}{m} + \binom{k-1}{a-\delta}$$

where $\delta = 0$ if $x_r$ is activatory and $\delta = 1$ if $x_r$ is inhibitory. If all the regulations including the self-regulation are inhibitory, the BMR corresponds to a constant-zero BF (The formula correctly yields a bias of 0, using the convention $\binom{\alpha}{\beta} = 0$ if $\beta < 0$). For $k > 2$, if the self-regulation is inhibitory and if there exists at least one activator among the other regulators, then $x_r$ is the only ineffective input to $f$ (see Property 2.4). For $k = 1$ and 2, the BF corresponding to a BMR with self-regulation is effective if and only if only all regulators are activatory. When $f$ is ineffective, after converting the $f$ to an EF, the bias will be $\frac{1}{2}P^k_{BMR}$. □

**Property 2.7.** A BF which is equal to a BMR with $k$ regulators, all being external, is canalyzing if and only if all $k$ regulators are either activatory or all are inhibitory.

*Proof.* The output of a BF equal to a BMR is determined by the values of the signed sum $S = \sum_{j=1}^{k} \sigma_j x_j$ and $x_{k+1}$ ($k+1$ being the target node) using the following rule:

$$f(x_1, x_2, \ldots, x_k, x_{k+1}) = \begin{cases} 1, & \text{if } S > 0 \\ 0, & \text{if } S < 0 \\ x_{k+1}, & \text{if } S = 0 \end{cases}$$

We first prove that if all the regulators are either activatory or inhibitory, then the corresponding BF is canalyzing. If all the regulators are activatory (i.e., $\sigma_i = +1 \; \forall \; 1 \le i \le k$), then the signed sum is given by $S = \sum_{j=1}^{k} x_j$. In this case, if $x_1 = 1$ then regardless of the values of remaining inputs, we have $S > 0$, and hence the output is 1. Therefore, $x_1$ acts as a canalyzing variable with canalyzing input value 1 and canalyzed output value 1. Similarly, when all the regulators are inhibitory, setting any $x_i = 1$, regardless of the other inputs the signed sum is negative and hence the output is 0. Thus, each $x_i$ for $1 \leq i \leq k+1$ is a canalyzing input with canalyzing input value 1 and canalyzed output value 0.

Now we will show that if not all the regulators are of same type then the BF cannot be canalyzing. Suppose, $f$ is canalyzing but not all regulator signs are of the same type. So, $\exists\ p, q\ (p \neq q)$ such that $\sigma_p = +1$ and $\sigma_q = -1$. Since, $f$ is canalyzing then there must be some variable $x_r$ and $a \in \{0, 1\}$ such that $f(x_1, \ldots, x_{r-1}, a, x_{r+1}, \ldots x_{k+1}) = c$ for some fixed value of $c \in \{0, 1\}$. Now, let us consider different cases separately by fixing $x_r = a$ and comparing the outcomes for two distinct input configurations.

**Case 1:** $\sigma_r = +1$
**Subcase 1.1:** $a = 0$
**Input 1:** $x_i = 0\ \forall\ i \neq r$ and $x_{k+1} = 0$. Then, the output is 0.
**Input 2:** $x_i = 0\ \forall\ i \neq r$ and $x_{k+1} = 1$. Then, the output is 1.
**Subcase 1.2:** $a = 1$
**Input 1:** $x_i = 0\ \forall\ i \neq r$ and $x_{k+1} = 0$. Then, the output is 1.
**Input 2:** $x_q = 1$, $x_i = 0\ \forall\ i \neq q, r$ and $x_{k+1} = 0$. Then, the output is 0.

**Case 2:** $\sigma_r = -1$
**Subcase 2.1:** $a = 0$
**Input 1:** $x_i = 0\ \forall\ i \neq r$ and $x_{k+1} = 0$. Then, the output is 0.
**Input 2:** $x_i = 0\ \forall\ i \neq r$ and $x_{k+1} = 1$. Then, the output is 1.
**Subcase 2.2:** $a = 1$
**Input 1:** $x_i = 0\ \forall\ i \neq r$ and $x_{k+1} = 0$. Then, the output is 0.
**Input 2:** $x_p = 1$, $x_i = 0\ \forall\ i \neq p, r$ and $x_{k+1} = 1$. Then, the output is 1.

Thus, we observe that in every possible condition, a contradiction arises, implying that none of the variables $x_i$ for $1 \leq i \leq k$ can serve as a canalyzing variable. It is also straightforward to verify that $x_{k+1}$ cannot be canalyzing either. To illustrate, consider the following two input configurations: (1) $x_p = 1$ and $x_i = 0$ for all $i \neq p$, and (2) $x_q = 1$ and $x_i = 0$ for all $i \neq q$. For any fixed value of $x_{k+1}$, the first input configurations lead to the output 1 whereas the second input leads to the output 0. This demonstrates that a BF equal to a BMR with $k$ external regulators (i.e., without a self-regulation) is canalyzing if and only if all $k$ regulators are either activatory or all are inhibitory. □

**Property 2.8.** A BF that is equal to a BMR with $k$ regulators including a self-regulation is canalyzing if and only if at most one regulator is activatory or at most one regulator is inhibitory.

*Proof.* We first show that if there is at most one inhibitory regulator or at most one activatory regulator, then the BF $f$ is canalyzing. Let $k$ regulators be $x_1, x_2, \ldots, x_k$. Without loss of generality, assume that $x_k$ is the self-regulatory input.

**Case 1 (At most one activator)**: If there is no activator then $f$ is constant zero function (See Property 2.4(c)) and hence canalyzing. When there is exactly one activator $x_r$, consider the case $x_r = 0$ as it will be canalyzing. Indeed,

$$S = \sum_{i \neq r} \sigma_i x_i = -\sum_{i \neq r} x_i \leq 0$$

Clearly, if $S < 0$ the output is 0; if instead $S = 0$, then necessarily $x_i = 0$ for all $i$ and in particular $x_k = 0$, so the output is also 0. Since all cases for which $x_r = 0$ lead to a output of 0, $x_r$ is a canalyzing variable.

**Case 2 (At most one inhibitor)**: If there is no inhibitor then the signed sum is $\sum_{i=1}^{k} x_i$. So if any $x_i = 1$, the signed sum is positive and hence the output is fixed to 1. If there is exactly one inhibitor $x_r$ and $r \neq k$, then if we fix $x_k = 1$, the signed sum becomes

$$S = -x_r + \sum_{i \neq r,k} x_i + 1 \geq 0$$

in which case the output is always 1 since $x_k = 1$. Now, if instead $k = r$, then $x_k$ is ineffective (see Property 2.4(b)) and the signed sum becomes $S = \sum_{i=1}^{k-1} x_i$. Hence for any $x_i$ $(1 \leq i < k)$ fixed to 1, the signed sum is positive and hence the output is fixed to 1.

We have shown that if there is at most one activator or at most one inhibitor among the regulators, the BF $f$ equal to the BMR is canalyzing. We now prove the converse. Specifically, we show that if there exist at least two activatory and at least two inhibitory regulators, then $f$ cannot be canalyzing. Suppose $x_t$ is a canalyzing variable for $f$ with the canalyzing input value $a \in \{0, 1\}$ where $t < k$. Since, there exists at least two activators and at least two inhibitors $\exists\ p, q \neq t$ such that $\sigma_p = +1, \sigma_q = -1$.

**Input case 1**: $x_t = a, x_p = 1, x_{i \notin \{t,p\}} = 0, x_k = 1$. The signed sum $S = \sigma_t a + 1 \geq 0$. So the output is always 1 as $x_k = 1$.

**Input case 2**: $x_t = a, x_q = 1, x_{i \notin \{t,q\}} = 0, x_k = 0$. The signed sum $S = \sigma_t a - 1 \leq 0$. So the output is always 0 as $x_k = 0$.

As a result, none of the $x_t$ can be a canalyzing variables for $t < k$. Now we will show that $x_k$ cannot be canalyzing either. Let us fix $x_k = a \in \{0, 1\}$. Choosing $p, q < k$ such that $\sigma_p = +1, \sigma_q = -1$ allows us to select two input combinations such that one leads to the output 1 whereas the other leads to the output 0 while keeping $x_k = a$ fixed.

**Input case 1**: $x_p = 1$ and $x_i = 0\ \forall\ i \neq p, k$. Then $S = 1 + \sigma_k a \geq 0$. Now if $1 + \sigma_k a > 0$, the output is 1. $S$ can be equal to 0 only when $\sigma_k = -1, a = 1$. In that case too, the output is 1 since $x_k = a = 1$.

**Input case 2**: Here we further break it down into two subcases:
**Subcase 1** ($\sigma_k = 1$): Choose $m \neq q$ such that $\sigma_m = -1$. Use the input: $x_k = a, x_q = x_m = 1$, all other $x_i = 0$. Then, $S = -2 + a < 0$. So, the output is always 0.
**Subcase 2** ($\sigma_k = -1$): $x_k = a, x_q = 1$, all other $x_i = 0$. Then, $S = -1 - a < 0$. The output is again 0.

So, this proves that a BF which is equal to a BMR with $k$ regulators and contains a self-regulation is canalyzing if and only if there is at most one activator or at most one inhibitor. ☐

**Property 2.9.** Let $f_1$ and $f_2$ denote the BFs corresponding to a $k$-input IMR and a $k$-input BMR, respectively, satisfying a given sign assignment of the regulators. Then, the following statements hold: (a) If $k$ is even, then $f_1 = f_2$ if and only if the number of activatory and inhibitory regulators are equal. (b) If $k$ is odd and $k > 1$, then $f_1$ and $f_2$ are never equal.

*Proof.* (a) First, consider the case when $k$ is even and one of the $k$ regulators is the self-regulator. Without loss of generality, assume that it is the $k^{\text{th}}$ regulator that is the target node itself. Let $S_I$ and $S_B$ denote the signed sums corresponding to IMR and BMR, respectively.

$$S_I(x') = \sum_{i=1}^{k} \sigma_i x'_i; \quad S_B(x) = \sum_{i=1}^{k} \sigma_i x_i$$

where $x = (x_1, x_2, \ldots, x_k)$, $x' = (x'_1, x'_2, \ldots x'_k)$ and $x'_i = 2x_i - 1$. Now for IMR (respectively, BMR), the output is active if $S_I > 0$ (similarly, $S_B > 0$) or $S_I = 0$ and $x'_k = +1$ (respectively, $S_B = 0$ and $x_k = 1$). Now, $S_I(x') = \sum_{i=1}^{k} \sigma_i x'_i = \sum_{i=1}^{k} \sigma_i(2x_i - 1) = 2S_B(x) - \Sigma$, where, $\Sigma = \sum_{i=1}^{k} \sigma_i$. Thus, we have the following three conditions of equivalency: (i) $S_I > 0 \iff S_B > \frac{\Sigma}{2}$; (ii) $S_I = 0 \iff S_B = \frac{\Sigma}{2}$; and (iii) $x'_k = +1 \iff x_k = 1$ (since, $x'_k = 2x_k - 1$).

Now if the number of activators and inhibitors are equal, then $\Sigma = 0$. Then, using the three equivalency conditions, $f_1$ and $f_2$ are either both active or both inactive for every input $x'$ to IMR and for the corresponding $x$ input to BMR, i.e., $f_1 = f_2$ up to the mapping between $x$ and $x'$.

We can now prove the converse. We will show that if the number of activators and inhibitors are unequal, i.e., if $\Sigma \neq 0$, then there exists $x'$ such that $f_1(x') \neq f_2(x)$. To proceed, we note that since $k$ is even, $\Sigma$ is an even integer.

**Case 1** ($\Sigma > 0$): Since $\Sigma$ is positive, the number of activators is strictly greater than 1 as $k \geq 2$. So, there exists $j \neq k$ such that $\sigma_j = +1$. For $f_2$, consider the input $x$ where $x_j = 1$ while for all other $i \neq j$, $x_i = 0$. Then, $S_B(x) = 1 > 0$ and hence the output is 'on' for $f_2$. Now, from the equivalency condition, the output for $f_1$ will be 'on' if $S_B(x) > \frac{\Sigma}{2}$ or $S_B(x) = \frac{\Sigma}{2}$ and $x_k = 1$. But none of these conditions are satisfied since $\frac{\Sigma}{2} \geq 1$ and $x_k = 0$ and as a result $f_1$ and $f_2$ cannot be equal.

**Case 2** ($\Sigma < 0$): Since $\Sigma$ is a negative even integer, $\frac{\Sigma}{2} \leq -1$. Now, for $f_2$ consider the input $x$ such that $x_i = 0$ for all $i$. Then, $S_B(x) = 0$ and $x_k = 0$ and hence the output would be 0 ('off'). Furthermore, since $S_B = 0$ and

$0 > -1 \geq \frac{\Sigma}{2}$, for $f_1$ the output will be +1. So again we see that $f_1$ and $f_2$ are not equal.

Exactly the same arguments can be applied if the target is not an explicit regulator. This proves the first part that if $k$ is even then $f_1 = f_2$ iff the number of activators and inhibitors are equal for a given sign assignment.

(b) Now we consider the case when $k$ is odd. If the target is not an explicit regulator, then $f_1$ corresponds to a $k$-input BF whereas $f_2$ corresponds to a $(k+1)$-input BF. Hence, in this case $f_1 \neq f_2$. However, we now examine the possibility of $f_1 = f_2$ when the target is an explicit regulator. We can demonstrate using a counter-example that $f_1$ can never be equal to $f_2$ in this case either. Since $k$ is odd, the signed sum can never be zero, leading us to consider the two cases: (i) $\Sigma > 0$ and (ii) $\Sigma < 0$.

**Case 1** ($\Sigma > 0$):

**Subcase 1.1** ($\Sigma = 1$): Then there are $\frac{(k+1)}{2}$ activators and $\frac{(k-1)}{2}$ inhibitors. We turn all the inhibitors on and $\frac{(k-1)}{2}$ activators on, ensuring $x_k = 1$. We keep all other regulators off. Then $S_B = 0$ (since $k \geq 3$) and since $x_k = 1$ the output is active. However, $S_I = 2S_B - \Sigma = -1 < 0$. So, the output is inactive. Hence, $f_1 \neq f_2$.

**Subcase 1.2** ($\Sigma \geq 3$): Since $\Sigma$ is positive and $k$ is odd, there exists at least $\frac{(k+1)}{2}$ activators, i.e., at least 2 activators because $k \geq 3$. Thus, there exists $j \neq k$ such that $\sigma_j = +1$. For $f_2$, consider an input $x$ such that $x_j = 1$ and $x_i = 0$ for all $i \neq j$. Then $S_B(x) = 1$ and hence the output is 'active'. But, $S_I(x') = 2.1 - \Sigma \leq 2 - 3 = -1 < 0$. So, the output is inactive and hence we can conclude $f_1 \neq f_2$

**Case 2** ($\Sigma < 0$): We consider an input $x$ such that $x_i = 0$ for all $i$. Then $S_B(x) = 0$ and since $x_k = 0$, the output is inactive. Now, $S_I(x') = 2.0 - \Sigma = -\Sigma > 0$ since $\Sigma < 0$. Hence, the output will be active. Thus, $f_1 \neq f_2$

This proves that if $k$ is odd and $k > 1$, then $f_1$ and $f_2$ can never be equal, regardless of the sign assignments. However, when $k = 1$ and the self-regulation is activatory, then $f_1$ and $f_2$ are equal. This is the only situation at odd $k$ for which $f_1 = f_2$. ☐

**Property 2.10.** A $k$-input BF which is equal to an IMR has the maximum average sensitivity among all $k$-input unate functions (UFs).

*Proof.* A result in [7] suggests that for any non-constant monotone BF $f$ with $k$ inputs, the average sensitivity $\overline{s}(f)$ has the following bounds:

$$\frac{k}{2^k} \leq \overline{s}(f) \leq \binom{k}{\lfloor k/2 \rfloor} \cdot \frac{\lceil k/2 \rceil}{2^{k-1}}$$

The upper bound is tight for the function 'MAJORITY'. In Boolean logic a majority function outputs 'False' when half or more inputs are 'False', and 'True' otherwise. The Majority rule exactly corresponds to the IMR when all regulatory inputs are activatory (i.e., have positive signs) with the exception that in case of a tie (i.e., when the number of 'True' and 'False' inputs are equal) the majority function outputs 'False'. However, in the IMR rule, a tie (i.e., when the signed sum equals zero) results in the output being determined by the state of the target node itself. IMR generalizes the Majority functions by allowing all possible assignments of regulatory signs. Monotone functions form a specific subset of the unate functions (UFs). Any UF can be transformed into a monotone function by appropriately negating some of its inputs [8]. Since the average sensitivity remains invariant under input negation, the maximum value of $\overline{s}(f)$ over the space of all UFs will be the same as that over all monotone functions and is achieved by the IMRs. It is important to note that the distinction in the tie-breaking scenarios for the two cases does not affect the value of the average sensitivity [7]. ☐

**Corollary 2.2.** For any odd $k$, a $k$-input BF equal to an IMR has the same average sensitivity as a $(k+1)$-input BF equal to an IMR.

*Proof.* Since $k$ is odd, introduce $m$ such that $k = 2m + 1$. The average sensitivity of the BF with $k$ inputs that is equal to an IMR is given by

$$\overline{s}(f_k) = \binom{k}{\lfloor k/2 \rfloor} \cdot \frac{\lceil k/2 \rceil}{2^{k-1}}$$

Noting that $\lfloor k/2 \rfloor = m$ and $\lceil k/2 \rceil = m + 1$, we have

$$\overline{s}(f_k) = \frac{\binom{2m+1}{m} \cdot (m+1)}{2^{2m}}$$

Thus the average senstivity of the BF with $(k+1)$ inputs that is equal to an IMR is given by

$$\overline{s}(f_{k+1}) = \frac{\binom{2m+2}{m+1} \cdot (m+1)}{2^{2m+1}} = \frac{2m+2}{m+1} \cdot \frac{\binom{2m+1}{m} \cdot (m+1)}{2^{2m+1}} = \frac{\binom{2m+1}{m} \cdot (m+1)}{2^{2m}} = \overline{s}(f_k)$$

which proves the statement. □

### SUPPLEMENTARY TABLES

TABLE S1. **Numbers of various subtypes of the EThFs in the theoretical space.** Here $k$ corresponds to the number of effective inputs to a BF. The columns under the grouping 'Number of BFs' report the total count of BFs in theory belonging to the classes EThF, NCF, IMR, BMR, non-NCF BMR, NCF BMR and non-BMR NCF, across varying values of $k$.

| $k$ | Number of BFs | | | | | | |
|---|---|---|---|---|---|---|---|
| | EThF | NCF | IMR | BMR | non-NCF BMR | NCF BMR | non-BMR NCF |
| 1 | 2 | 2 | 2 | 1 | 0 | 1 | 1 |
| 2 | 8 | 8 | 0 | 3 | 0 | 3 | 5 |
| 3 | 72 | 64 | 8 | 16 | 3 | 13 | 51 |
| 4 | 1536 | 736 | 32 | 43 | 38 | 5 | 731 |
| 5 | 86080 | 10624 | 32 | 106 | 100 | 6 | 10618 |
| 6 | 14487040 | 183936 | 192 | 249 | 242 | 7 | 183929 |
| 7 | 8274797440 | 3715072 | 128 | 568 | 560 | 8 | 3715064 |
| 8 | 17494930604032 | 85755372 | 1024 | 1271 | 1262 | 9 | 85755363 |
| 9 | 144222448789966828 | 2226939904 | 512 | 2806 | 2796 | 10 | 2226939894 |

TABLE S2. **Numbers and proportions of various subtypes of the EThFs in the BBM benchmark dataset.** Here $k$ corresponds to the number of effective inputs to a BF. The columns under the grouping 'Number of BFs' report the total count of BFs in the BBM dataset ('Total' column), along with the counts of BFs belonging to the classes EThF, NCF, IMR, BMR, non-NCF BMR, NCF BMR and non-BMR NCF, across varying values of $k$. The columns under grouping 'Fraction of subtypes within EThFs' presents the relative proportions of the different subtypes of EThF within EThFs for each $k$.

| $k$ | Number of BFs | | | | | | | | Fraction of subtypes within EThFs | | | | | |
|---|---|---|---|---|---|---|---|---|---|---|---|---|---|---|
| | Total | EThF | NCF | IMR | BMR | non-NCF BMR | NCF BMR | non-BMR NCF | NCF | IMR | BMR | non-NCF BMR | NCF BMR | non-BMR NCF |
| 1 | 2013 | 2013 | 2013 | 2013 | 1771 | 0 | 1771 | 242 | 1.0 | 1.0 | 0.88 | 0.0 | 0.88 | 0.12 |
| 2 | 1634 | 1634 | 1634 | 0 | 902 | 0 | 902 | 732 | 1.0 | 0.0 | 0.552 | 0.0 | 0.552 | 0.448 |
| 3 | 914 | 908 | 895 | 13 | 312 | 12 | 300 | 595 | 0.986 | 0.014 | 0.344 | 0.013 | 0.33 | 0.655 |
| 4 | 552 | 515 | 506 | 0 | 85 | 2 | 83 | 423 | 0.983 | 0.0 | 0.165 | 0.004 | 0.161 | 0.821 |
| 5 | 371 | 318 | 303 | 0 | 52 | 5 | 47 | 256 | 0.953 | 0.0 | 0.164 | 0.016 | 0.148 | 0.805 |
| 6 | 222 | 168 | 160 | 0 | 10 | 0 | 10 | 150 | 0.952 | 0.0 | 0.06 | 0.0 | 0.06 | 0.893 |
| 7 | 122 | 84 | 81 | 1 | 3 | 1 | 2 | 79 | 0.964 | 0.012 | 0.036 | 0.012 | 0.024 | 0.94 |
| 8 | 74 | 51 | 50 | 0 | 3 | 0 | 3 | 47 | 0.98 | 0.0 | 0.059 | 0.0 | 0.059 | 0.922 |
| 9 | 51 | 31 | 29 | 0 | 1 | 0 | 1 | 28 | 0.935 | 0.0 | 0.032 | 0.0 | 0.032 | 0.903 |

TABLE S3. **Numbers and proportions of various subtypes of the EThFs in the MCBF dataset.** Here $k$ corresponds to the number of effective inputs to a BF. The columns under the grouping 'Number of BFs' report the total count of BFs in the MCBF dataset ('Total' column), along with the counts of BFs belonging to the classes EThF, NCF, IMR, BMR, non-NCF BMR, NCF BMR and non-BMR NCF, across varying values of $k$. The columns under grouping 'Fraction of subtypes within EThFs' presents the relative proportions of the different subtypes of EThF within EThFs for each $k$.

| $k$ | Number of BFs | | | | | | | | Fraction of subtypes within EThFs | | | | | |
|---|---|---|---|---|---|---|---|---|---|---|---|---|---|---|
| | Total | EThF | NCF | IMR | BMR | non-NCF BMR | NCF BMR | non-BMR NCF | NCF | IMR | BMR | non-NCF BMR | NCF BMR | non-BMR NCF |
| 1 | 953 | 953 | 953 | 953 | 812 | 0 | 812 | 141 | 1.0 | 1.0 | 0.852 | 0.0 | 0.852 | 0.148 |
| 2 | 689 | 689 | 689 | 0 | 460 | 0 | 460 | 229 | 1.0 | 0.0 | 0.668 | 0.0 | 0.668 | 0.332 |
| 3 | 402 | 401 | 388 | 13 | 140 | 12 | 128 | 260 | 0.968 | 0.032 | 0.349 | 0.03 | 0.319 | 0.648 |
| 4 | 259 | 244 | 238 | 0 | 52 | 3 | 49 | 189 | 0.975 | 0.0 | 0.213 | 0.012 | 0.201 | 0.775 |
| 5 | 155 | 135 | 125 | 0 | 25 | 4 | 21 | 104 | 0.926 | 0.0 | 0.185 | 0.03 | 0.156 | 0.77 |
| 6 | 99 | 76 | 67 | 0 | 3 | 0 | 3 | 64 | 0.882 | 0.0 | 0.039 | 0.0 | 0.039 | 0.842 |
| 7 | 49 | 35 | 35 | 0 | 0 | 0 | 0 | 35 | 1.0 | 0.0 | 0.0 | 0.0 | 0.0 | 1.0 |
| 8 | 45 | 27 | 27 | 0 | 1 | 0 | 1 | 26 | 1.0 | 0.0 | 0.037 | 0.0 | 0.037 | 0.963 |
| 9 | 19 | 7 | 7 | 0 | 1 | 0 | 1 | 6 | 1.0 | 0.0 | 0.143 | 0.0 | 0.143 | 0.857 |

TABLE S4. **Numbers and proportions of various subtypes of the EThFs in the Harris dataset.** Here $k$ corresponds to the number of effective inputs to a BF. The columns under the grouping 'Number of BFs' report the total count of BFs in the Harris dataset ('Total' column), along with the counts of BFs belonging to the classes EThF, NCF, IMR, BMR, non-NCF BMR, NCF BMR and non-BMR NCF, across varying values of $k$. The columns under grouping 'Fraction of subtypes within EThFs' presents the relative proportions of the different subtypes of EThF within EThFs for each $k$.

| $k$ | Number of BFs | | | | | | | | Fraction of subtypes within EThFs | | | | | |
|---|---|---|---|---|---|---|---|---|---|---|---|---|---|---|
| | Total | EThF | NCF | IMR | BMR | non-NCF BMR | NCF BMR | non-BMR NCF | NCF | IMR | BMR | non-NCF BMR | NCF BMR | non-BMR NCF |
| 1 | 2 | 2 | 2 | 2 | 2 | 0 | 2 | 0 | 1.0 | 1 | 1.0 | 0 | 1.0 | 0.0 |
| 2 | 9 | 9 | 9 | 0 | 3 | 0 | 3 | 6 | 1.0 | 0 | 0.333 | 0 | 0.333 | 0.667 |
| 3 | 71 | 71 | 71 | 0 | 8 | 0 | 8 | 63 | 1.0 | 0 | 0.113 | 0 | 0.113 | 0.887 |
| 4 | 38 | 36 | 35 | 0 | 0 | 0 | 0 | 35 | 0.972 | 0 | 0.0 | 0 | 0.0 | 0.972 |
| 5 | 19 | 16 | 16 | 0 | 0 | 0 | 0 | 16 | 1.0 | 0 | 0.0 | 0 | 0.0 | 1.0 |

TABLE S5. **Relative enrichment ratios ($E_R$) and statistical significance ($p$-values) for EThF subtypes in the BBM benchmark dataset.** Here, $k$ is the number of effective inputs per BF. Under '$E_R$ of subtypes within EThFs', the columns 'NCF', 'IMR' and 'BMR' display the relative enrichment ratios ($E_R$) for each subtype at given $k$ (see Main text, Section 2.4). $E_R > 1$ indicates enrichment of the corresponding subtype within EThFs in the BBM dataset. No entry appears for subtype IMR at $k = 2$ because no 2-input EF is equal to such an IMR. Under '$p$-values associated with $E_R$ of different subtypes', the same columns report the significance of each enrichment (with $p < 0.05$ indicating statistical significance). These $p$-values have been used to annotate Fig. 2(c) in Main text. For $k = 1$, the subtypes NCF and IMR coincide with EThF, and for $k = 2$, NCF coincides with EThF while IMR is absent; accordingly, $p$-values are not shown in these cases.

| $k$ | $E_R$ of subtypes within EThFs | | | $p$-values associated with $E_R$ of different subtypes | | |
|---|---|---|---|---|---|---|
| | NCF | IMR | BMR | NCF | IMR | BMR |
| 1 | 1 | 1 | 1.76 | - | - | $4.85 \times 10^{-287}$ |
| 2 | 1 | - | 1.47 | - | - | $9.59 \times 10^{-48}$ |
| 3 | 1.11 | 0.13 | 1.55 | $3.09 \times 10^{-30}$ | 1 | $4.47 \times 10^{-17}$ |
| 4 | 2.05 | 0 | 5.90 | $3.98 \times 10^{-146}$ | 1 | $4.27 \times 10^{-39}$ |
| 5 | 7.72 | 0 | $1.33 \times 10^{2}$ | $1.29 \times 10^{-251}$ | 1 | $7.3 \times 10^{-92}$ |
| 6 | 75.01 | 0 | $3.46 \times 10^{3}$ | $4.67 \times 10^{-291}$ | 1 | $8.43 \times 10^{-33}$ |
| 7 | $2.15 \times 10^{3}$ | $7.7 \times 10^{5}$ | $5.2 \times 10^{5}$ | $6.42 \times 10^{-267}$ | $1.3 \times 10^{-6}$ | $3.08 \times 10^{-17}$ |
| 8 | $2.0 \times 10^{5}$ | 0 | $8.1 \times 10^{8}$ | $1.68 \times 10^{-264}$ | 1 | $7.99 \times 10^{-27}$ |
| 9 | $6.06 \times 10^{7}$ | 0 | $1.66 \times 10^{12}$ | $1.38 \times 10^{-224}$ | 1 | $6.03 \times 10^{-13}$ |

TABLE S6. **Relative enrichment ratios ($E_R$) and statistical significance ($p$-values) for the EThF subtypes 'non-BMR NCF', 'NCF BMR' and 'non-NCF BMR' in the BBM benchmark dataset.** Here, $k$ is the number of effective inputs per BF. A, B and C denote the subtypes non-BMR NCF, NCF BMR and non-NCF BMR, respectively. Under '$E_R$ of subtypes within EThFs', the columns display the relative enrichment ratios ($E_R$) for each subtype at given $k$ (see Main text, Section 2.4). $E_R > 1$ indicates enrichment of the corresponding subtype within EThFs in the BBM dataset. No entry appears for column 'C' at $k = 1$ and 2 because there are no EFs of the type 'non-NCF BMR' for $k = 1$ and 2. Under '$p$-values associated with $E_R$ of different subtypes and their order', the same columns report the significance of each enrichment. These $p$-values have been used to annotate Fig. 2(d) in Main text. As for $k = 1$ and 2, no EF belongs to the non-NCF BMR category, so the corresponding $p$-values are omitted. The 'Order' column ranks subtypes by increasing $p$-value; larger $p$-values indicate weaker statistical significance for enrichment.

| $k$ | $E_R$ of subtypes within EThFs | | | $p$-values associated with $E_R$ of different subtypes and their order | | | |
|---|---|---|---|---|---|---|---|
| | A | B | C | A | B | C | Order |
| 1 | 0.24 | 1.76 | - | 1 | $4.85 \times 10^{-287}$ | - | B < A |
| 2 | 0.72 | 1.47 | - | 1 | $9.59 \times 10^{-48}$ | - | B < A |
| 3 | 0.93 | 1.83 | 0.32 | 1.00 | $1.99 \times 10^{-27}$ | 1 | B < A < C |
| 4 | 1.73 | 49.51 | 0.16 | $3.2 \times 10^{-59}$ | $2.44 \times 10^{-110}$ | 1.00 | B < A < C |
| 5 | 6.53 | $2.12 \times 10^{3}$ | 13.53 | $4.87 \times 10^{-170}$ | $1.87 \times 10^{-139}$ | $4.11 \times 10^{-5}$ | A < B < C |
| 6 | 70.33 | $1.23 \times 10^{5}$ | 0 | $1.95 \times 10^{-261}$ | $2.61 \times 10^{-48}$ | 1 | A < B < C |
| 7 | $2.09 \times 10^{3}$ | $2.46 \times 10^{7}$ | $1.76 \times 10^{5}$ | $1.03 \times 10^{-257}$ | $3.26 \times 10^{-15}$ | $5.68 \times 10^{-6}$ | A < B < C |
| 8 | $1.88 \times 10^{5}$ | $1.14 \times 10^{11}$ | 0 | $6.98 \times 10^{-245}$ | $2.84 \times 10^{-33}$ | 1 | A < B < C |
| 9 | $5.85 \times 10^{7}$ | $4.65 \times 10^{14}$ | 0 | $8.62 \times 10^{-216}$ | $2.15 \times 10^{-15}$ | 1 | A < B < C |

TABLE S7. **Relative enrichment ratios ($E_R$) and statistical significance ($p$-values) for EThF subtypes in the MCBF dataset.** Here, $k$ is the number of effective inputs per BF. Under '$E_R$ of subtypes within EThFs', the columns 'NCF', 'IMR' and 'BMR' display the relative enrichment ratios ($E_R$) for each subtype at given $k$ (see Main text, Section 2.4). $E_R > 1$ indicates enrichment of the corresponding subtype within EThFs in the MCBF dataset. No entry appears for subtype IMR at $k = 2$ because no 2-input EF is equal to such an IMR. Under '$p$-values associated with $E_R$ of different subtypes', the same columns report the significance of each enrichment (with $p < 0.05$ indicating statistical significance). These $p$-values have been used to annotate SI Fig. S2(c). For $k = 1$, the subtypes NCF and IMR coincide with EThF, and for $k = 2$, NCF coincides with EThF while IMR is absent; accordingly, $p$-values are not shown in these cases.

| $k$ | $E_R$ of subtypes within EThFs | | | $p$-values associated with $E_R$ of different subtypes | | |
|---|---|---|---|---|---|---|
| | NCF | IMR | BMR | NCF | IMR | BMR |
| 1 | 1 | 1 | 1.70 | - | - | $1.73 \times 10^{-115}$ |
| 2 | 1 | - | 1.78 | - | - | $1.72 \times 10^{-54}$ |
| 3 | 1.09 | 0.29 | 1.57 | $6.91 \times 10^{-9}$ | 1 | $4.63 \times 10^{-9}$ |
| 4 | 2.04 | 0 | 7.61 | $5.08 \times 10^{-67}$ | 1 | $4.16 \times 10^{-30}$ |
| 5 | 7.50 | 0 | $1.5 \times 10^{2}$ | $2.83 \times 10^{-100}$ | 1 | $1.74 \times 10^{-46}$ |
| 6 | 69.43 | 0 | $2.3 \times 10^{3}$ | $1.13 \times 10^{-116}$ | 1 | $3.57 \times 10^{-10}$ |
| 7 | $2.23 \times 10^{3}$ | 0 | 0 | $6.72 \times 10^{-118}$ | 1 | 1 |
| 8 | $2.04 \times 10^{5}$ | 0 | $5.1 \times 10^{8}$ | $4.36 \times 10^{-144}$ | 1 | $1.96 \times 10^{-9}$ |
| 9 | $6.48 \times 10^{7}$ | 0 | $7.34 \times 10^{12}$ | $2.09 \times 10^{-55}$ | 1 | $1.36 \times 10^{-13}$ |

TABLE S8. **Relative enrichment ratios ($E_R$) and statistical significance ($p$-values) for the EThF subtypes 'non-BMR NCF', 'NCF BMR' and 'non-NCF BMR' in the MCBF dataset.** Here, $k$ is the number of effective inputs per BF. A, B and C denote the subtypes non-BMR NCF, NCF BMR and non-NCF BMR, respectively. Under '$E_R$ of subtypes within EThFs', the columns display the relative enrichment ratios ($E_R$) for each subtype at given $k$ (see Main text, Section 2.4). $E_R > 1$ indicates enrichment of the corresponding subtype within EThFs in the MCBF dataset. No entry appears for column 'C' at $k = 1$ and 2 because there are no EFs of the type 'non-NCF BMR' for $k = 1$ and 2. Under '$p$-values associated with $E_R$ of different subtypes and their order', the same columns report the significance of each enrichment. These $p$-values have been used to annotate SI Fig. S2(d) in Main text. As for $k = 1$ and 2, no EF belongs to the non-NCF BMR category, so the corresponding $p$-values are omitted. The 'Order' column ranks subtypes by increasing $p$-value; larger $p$-values indicate weaker statistical significance for enrichment.

| $k$ | $E_R$ of subtypes within EThFs | | | $p$-values associated with $E_R$ of different subtypes and their order | | | |
|---|---|---|---|---|---|---|---|
| | A | B | C | A | B | C | Order |
| 1 | 0.30 | 1.70 | - | 1 | $1.73 \times 10^{-115}$ | - | B < A |
| 2 | 0.53 | 1.78 | - | 1 | $1.72 \times 10^{-54}$ | - | B < A |
| 3 | 0.92 | 1.77 | 0.72 | 1.00 | $1.59 \times 10^{-11}$ | 0.91 | B < C < A |
| 4 | 1.63 | 61.69 | 0.50 | $1.23 \times 10^{-21}$ | $6.26 \times 10^{-71}$ | 0.94 | B < A < C |
| 5 | 6.25 | $2.23 \times 10^{3}$ | 25.51 | $1.69 \times 10^{-66}$ | $1.05 \times 10^{-63}$ | $2.13 \times 10^{-5}$ | A < B < C |
| 6 | 66.33 | $8.17 \times 10^{4}$ | 0 | $1.15 \times 10^{-108}$ | $7.93 \times 10^{-15}$ | 1 | A < B < C |
| 7 | $2.23 \times 10^{3}$ | 0 | 0 | $6.72 \times 10^{-118}$ | 1 | 1 | A < B = C |
| 8 | $1.96 \times 10^{5}$ | $7.2 \times 10^{10}$ | 0 | $2.4 \times 10^{-137}$ | $1.39 \times 10^{-11}$ | 1 | A < B < C |
| 9 | $5.55 \times 10^{7}$ | $2.06 \times 10^{15}$ | 0 | $9.49 \times 10^{-47}$ | $4.85 \times 10^{-16}$ | 1 | A < B < C |

TABLE S9. **Relative enrichment ratios ($E_R$) and statistical significance ($p$-values) for EThF subtypes in the Harris dataset.** Here, $k$ is the number of effective inputs per BF. Under '$E_R$ of subtypes within EThFs', the columns 'NCF', 'IMR' and 'BMR' display the relative enrichment ratios ($E_R$) for each subtype at given $k$ (see Main text, Section 2.4). $E_R > 1$ indicates enrichment of the corresponding subtype within EThFs in the Harris dataset. No entry appears for subtype IMR at $k = 2$ because no 2-input EF is equal to such an IMR. Under '$p$-values associated with $E_R$ of different subtypes', the same columns report the significance of each enrichment (with $p < 0.05$ indicating statistical significance). These $p$-values have been used to annotate SI Fig. S3(c). For $k = 1$, the subtypes NCF and IMR coincide with EThF, and for $k = 2$, NCF coincides with EThF while IMR is absent; accordingly, $p$-values are not shown in these cases.

| $k$ | $E_R$ of subtypes within EThFs | | | $p$-values associated with $E_R$ of different subtypes | | |
|---|---|---|---|---|---|---|
| | NCF | IMR | BMR | NCF | IMR | BMR |
| 1 | 1 | 1 | 2.00 | - | - | 0.50 |
| 2 | 1 | - | 1.33 | - | - | 0.61 |
| 3 | 1.12 | 0 | 1.29 | 0.44 | 1 | 0.48 |
| 4 | 1.90 | 0 | 0 | $3.96 \times 10^{-3}$ | 1 | 1 |
| 5 | 8.10 | 0 | 0 | $8.2 \times 10^{-10}$ | 1 | 1 |

TABLE S10. **Relative enrichment ratios ($E_R$) and statistical significance ($p$-values) for the EThF subtypes 'non-BMR NCF', 'NCF BMR' and 'non-NCF BMR' in the Harris dataset.** Here, $k$ is the number of effective inputs per BF. A, B and C denote the subtypes non-BMR NCF, NCF BMR and non-NCF BMR, respectively. Under '$E_R$ of subtypes within EThFs', the columns display the relative enrichment ratios ($E_R$) for each subtype at given $k$ (see Main text, Section 2.4). $E_R > 1$ indicates enrichment of the corresponding subtype within EThFs in the Harris dataset. No entry appears for column 'C' at $k = 1$ and 2 because there are no EFs of the type 'non-NCF BMR' for $k = 1$ and 2. Under '$p$-values associated with $E_R$ of different subtypes and their order', the same columns report the significance of each enrichment. These $p$-values have been used to annotate SI Fig. S3(d) in Main text. As for $k = 1$ and 2, no EF belongs to the non-NCF BMR category, so the corresponding $p$-values are omitted. The 'Order' column ranks subtypes by increasing $p$-value; larger $p$-values indicate weaker statistical significance for enrichment.

| $k$ | $E_R$ of subtypes within EThFs | | | $p$-values associated with $E_R$ of different subtypes and their order | | | |
|---|---|---|---|---|---|---|---|
| | A | B | C | A | B | C | Order |
| 1 | 0 | 2.00 | - | 1 | 0.25 | - | B < A |
| 2 | 1.07 | 0.89 | - | 0.55 | 0.72 | - | A < B |
| 3 | 1.25 | 0.62 | 0 | $2.87 \times 10^{-4}$ | 0.96 | 1 | A < B < C |
| 4 | 2.04 | 0 | 0 | $1.0 \times 10^{-10}$ | 1 | 1 | A < B = C |
| 5 | 8.11 | 0 | 0 | $2.87 \times 10^{-15}$ | 1 | 1 | A < B = C |

TABLE S11. **Summary of models following replacement of non-NCF BFs with NCFs.** Each row corresponds to a specific network, identified by its ID as given in the BBM dataset. Column 'N' gives the number of nodes in the network. and column 'M' indicates the number of nodes whose assigned BF in the published model is non-NCF. The column 'Number of replaced models' denotes the total number of candidate models obtained by replacing all non-NCFs with valid scNCFs, such that the resulting model still recovers the biological attractors. The column 'Number of models for analysis' refers to the subset of these models that were randomly sampled (if too large) or fully enumerated for further dynamical analysis. The next three columns report how many models exhibit attractor recovery scores that are higher than or equal to the published model and higher than the IMR- and BMR-type variants respectively. The final three columns show how many models yield JS distances (from the gold standard basin fraction distribution) that are lower than or equal to that of the published model and are lower than those for the IMR- and BMR-type variants, respectively.

| ID | N | M | Number of replaced models | Number of models considered for analysis | Number of models with $\mathrm{ARS}_{repl} \geq \mathrm{ARS}_{model}$ | Number of models with $\mathrm{ARS}_{repl} > \mathrm{ARS}_{\mathrm{IMR}}$ | Number of models with $\mathrm{ARS}_{repl} > \mathrm{ARS}_{\mathrm{BMR}}$ | Number of models with $D_{repl} \leq D_{model}$ | Number of models with $D_{repl} < D_{\mathrm{IMR}}$ | Number of models with $D_{repl} < D_{\mathrm{BMR}}$ |
|---|---|---|---|---|---|---|---|---|---|---|
| 61 | 26 | 2 | 686244 | 5000 | 1640 | 5000 | 5000 | 4064 | 5000 | 5000 |
| 69 | 22 | 2 | 96 | 96 | 96 | 96 | 96 | 96 | 96 | 96 |
| 95 | 10 | 4 | 2116 | 2116 | 576 | 2080 | 2116 | 313 | 2116 | 2116 |
| 212 | 16 | 1 | 48 | 48 | 9 | 48 | 48 | 4 | 48 | 48 |

### SUPPLEMENTARY FIGURES

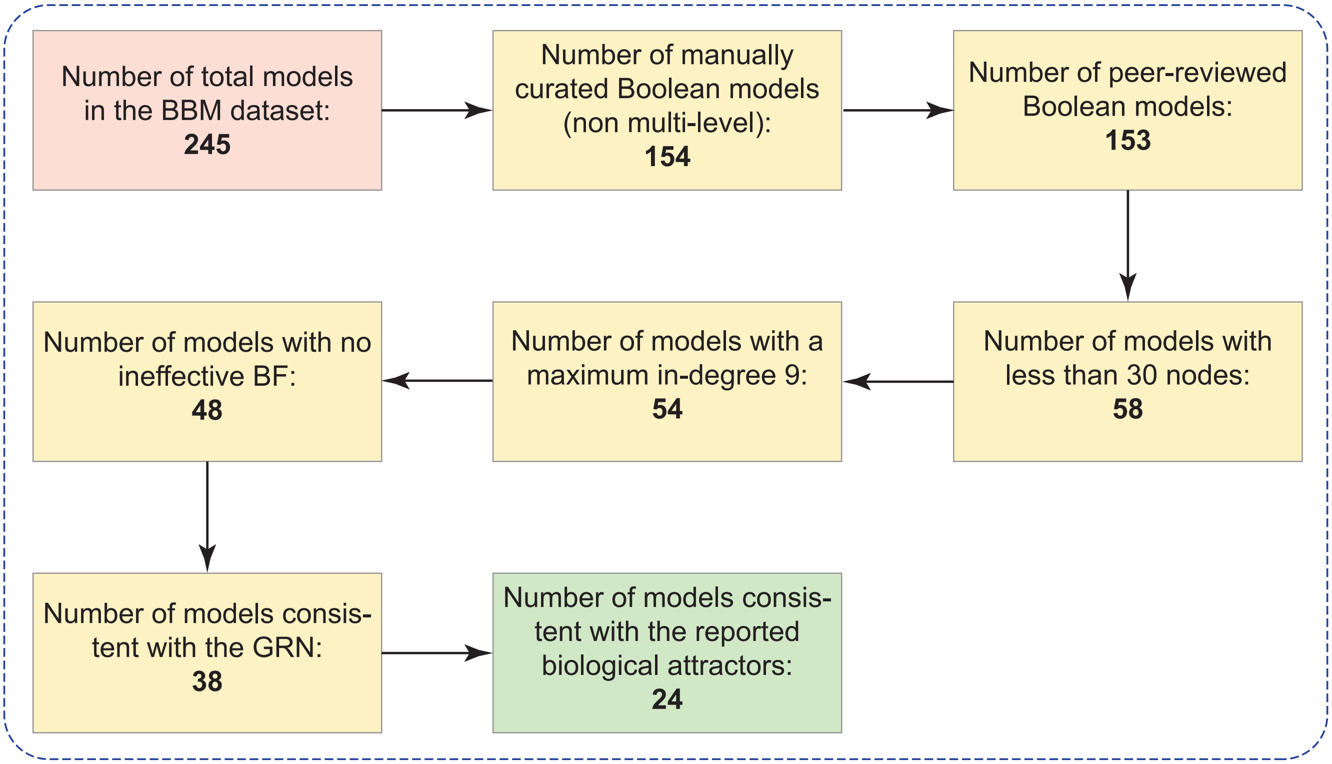


FIG. S1. **Flowchart of the multi-step filtration pipeline applied to the models in the BBM benchmark dataset.** Starting with 245 models from the BBM dataset, a series of filtration steps (as shown in the flowchart) resulted in a final set of 24 models selected for our detailed analysis of attractor recovery and associated basin sizes.

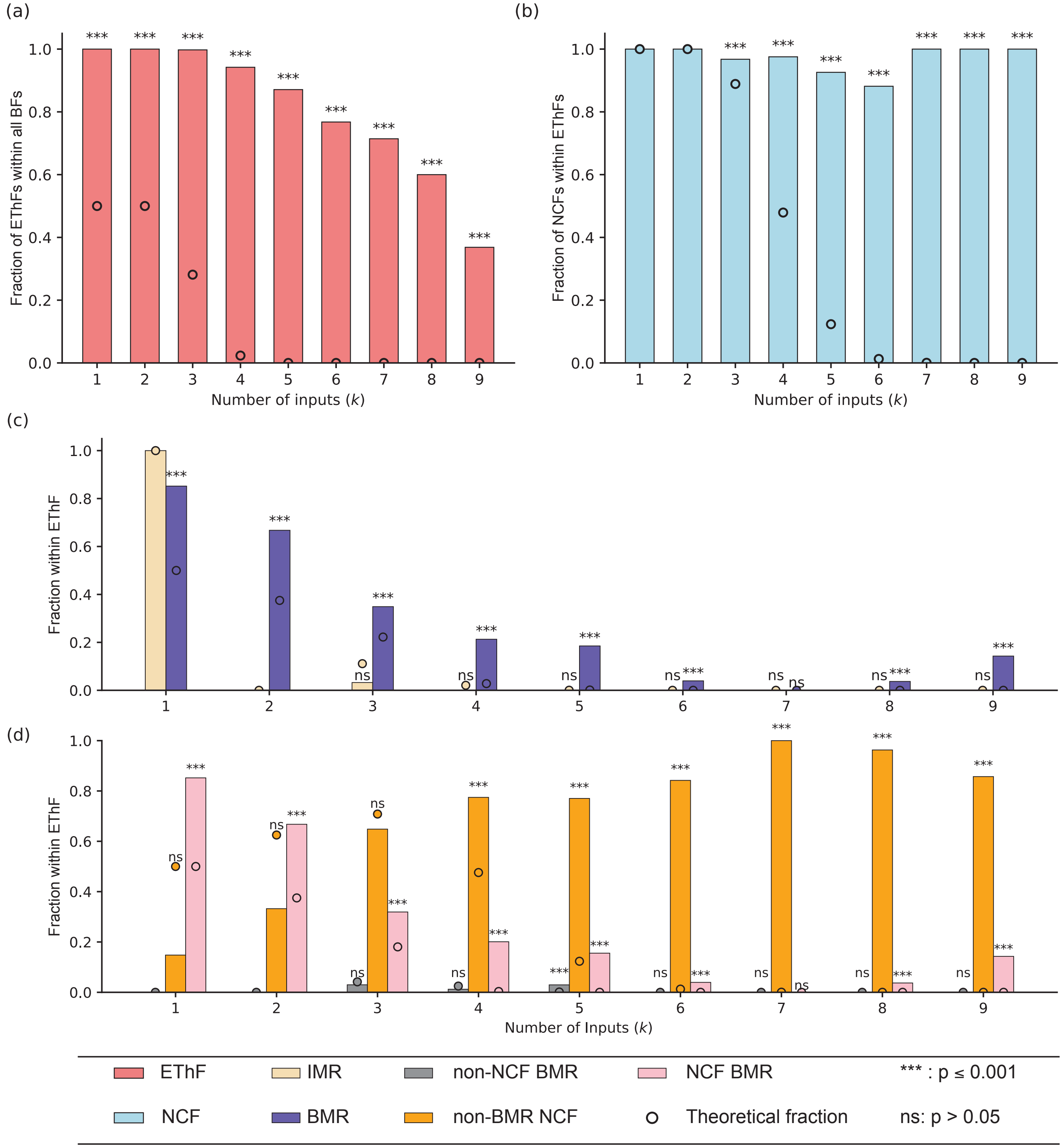


FIG. S2. **Fraction of various subtypes within an englobing type inside the MCBF dataset.** **(a)** The fraction of EThFs within all BFs for various number of inputs. **(b)** The fraction of NCFs within EThFs for various number of inputs. **(c)** The fraction of BFs that are equal to some IMR or BMR within the NCFs for various number of inputs. **(d)** The fractions of non-NCF BMRs, non-BMR NCFs and NCF BMRs within EThFs for various number of inputs. In all these plots, the theoretically expected values are represented by dots while the empirically obtained values from the MCBF dataset are shown via the color bars.

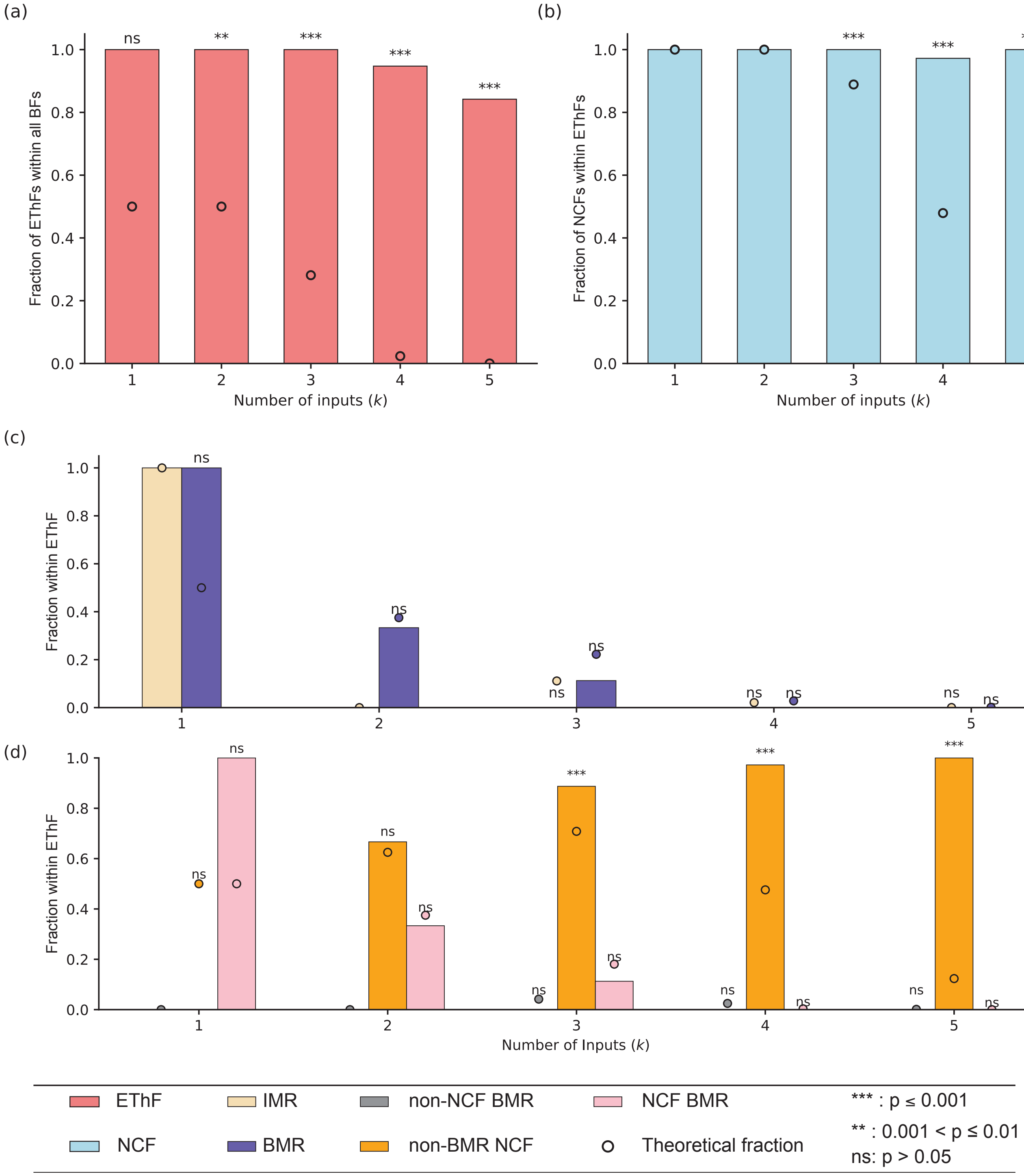


FIG. S3. **Fraction of various subtypes within an englobing type inside the Harris dataset.** **(a)** The fraction of EThFs within all BFs for various number of inputs. **(b)** The fraction of NCFs within EThFs for various number of inputs. **(c)** The fraction of BFs that are equal to some IMR or BMR within the NCFs for various number of inputs. **(d)** The fractions of non-NCF BMRs, non-BMR NCFs and NCF BMRs within EThFs for various number of inputs. In all these plots, the theoretically expected values are represented by dots while the empirically obtained values from the Harris dataset are shown via the color bars.

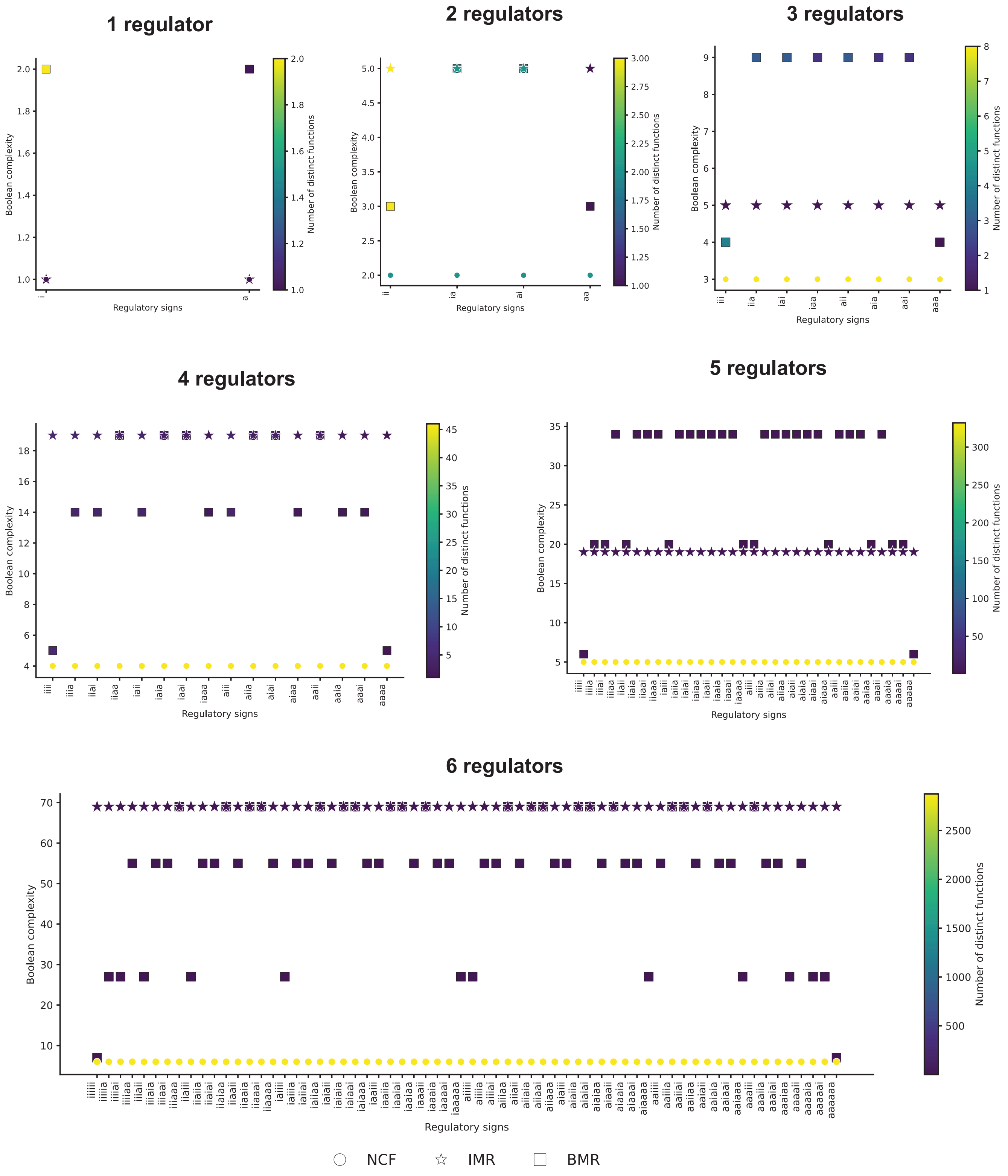


FIG. S4. **Boolean complexities for different sign combinations of the regulators**. Each of the subplots display the Boolean complexities of IMRs, BMRs and NCFs for different combinations of the signs of the regulators. In each subplot, for a given sign combination, the number of distinct functions that has a specific value of the Boolean complexity is shown using a color gradiant.

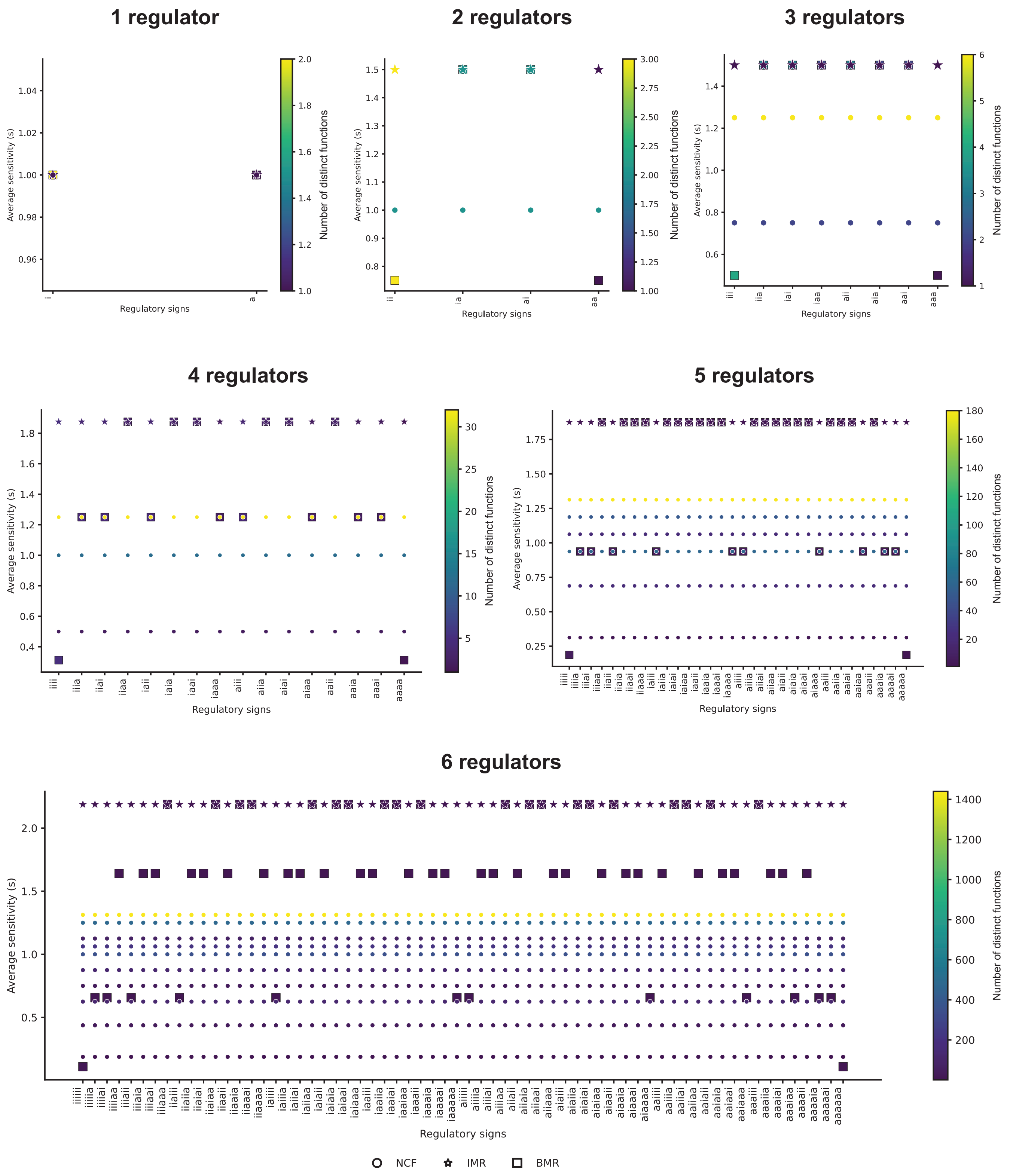


FIG. S5. **Average sensitivities for different sign combinations of the regulators**. Each of the subplots display the average sensitivities of IMRs, BMRs and NCFs for different combinations of the signs of the regulators. In each subplot, for a given sign combination, the number of distinct functions that has a specific value of the average sensitivity is shown using a color gradiant.

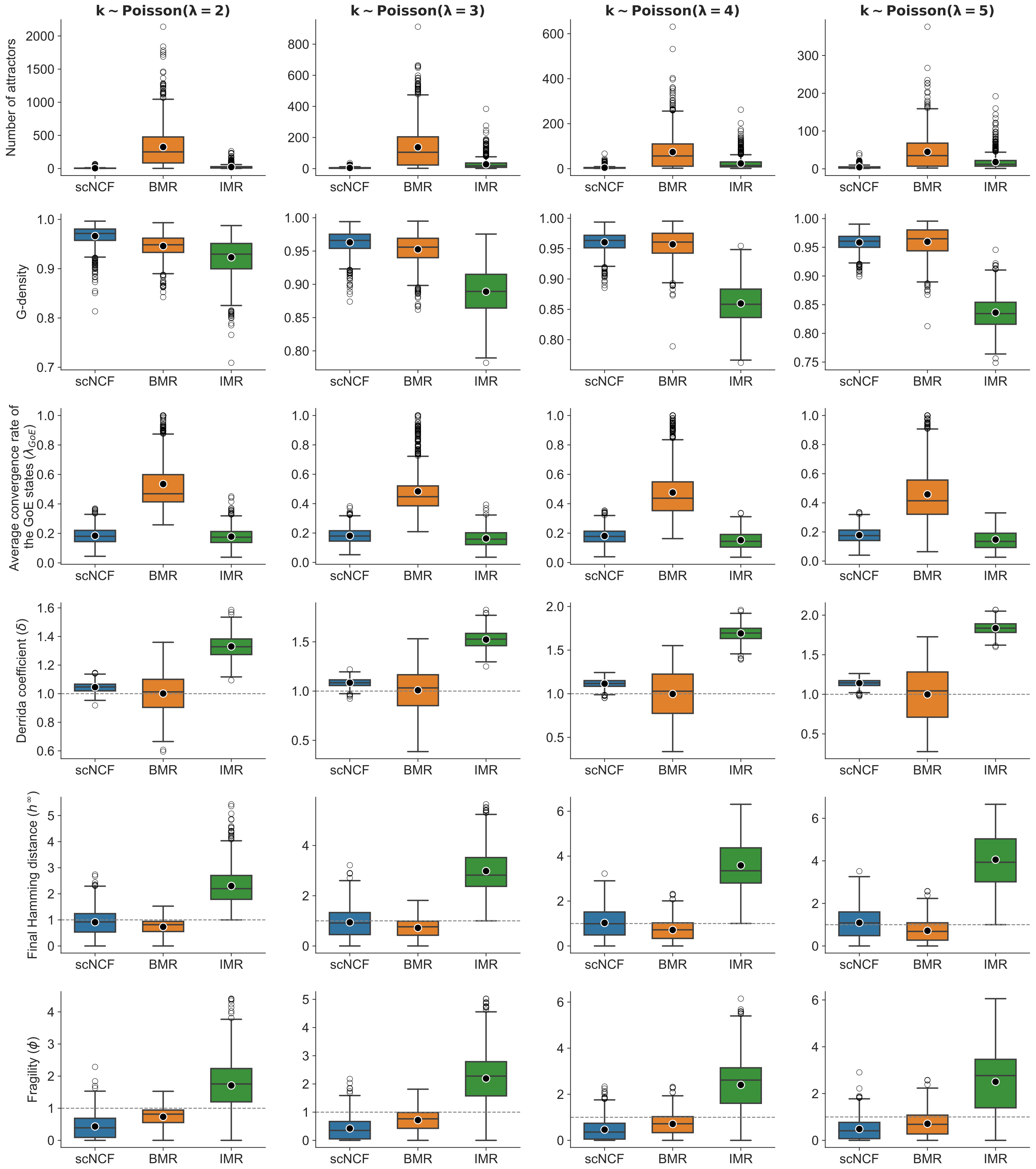


FIG. S6. **Distributions of stability measures for different subtypes of ThFs under the P-P network topology with $N = 16$.** Each row corresponds to one observable and each of the four columns represent different values of the mean in-degree parameter ($\lambda = 2, 3, 4, 5$). In each subplot, the three boxplots present the distributions of the observables across three subtypes of ThFs: scNCFs, BMRs and IMRs. Outliers in each distribution are shown as grey circles while the black dots outlined in white denote the mean of each group.

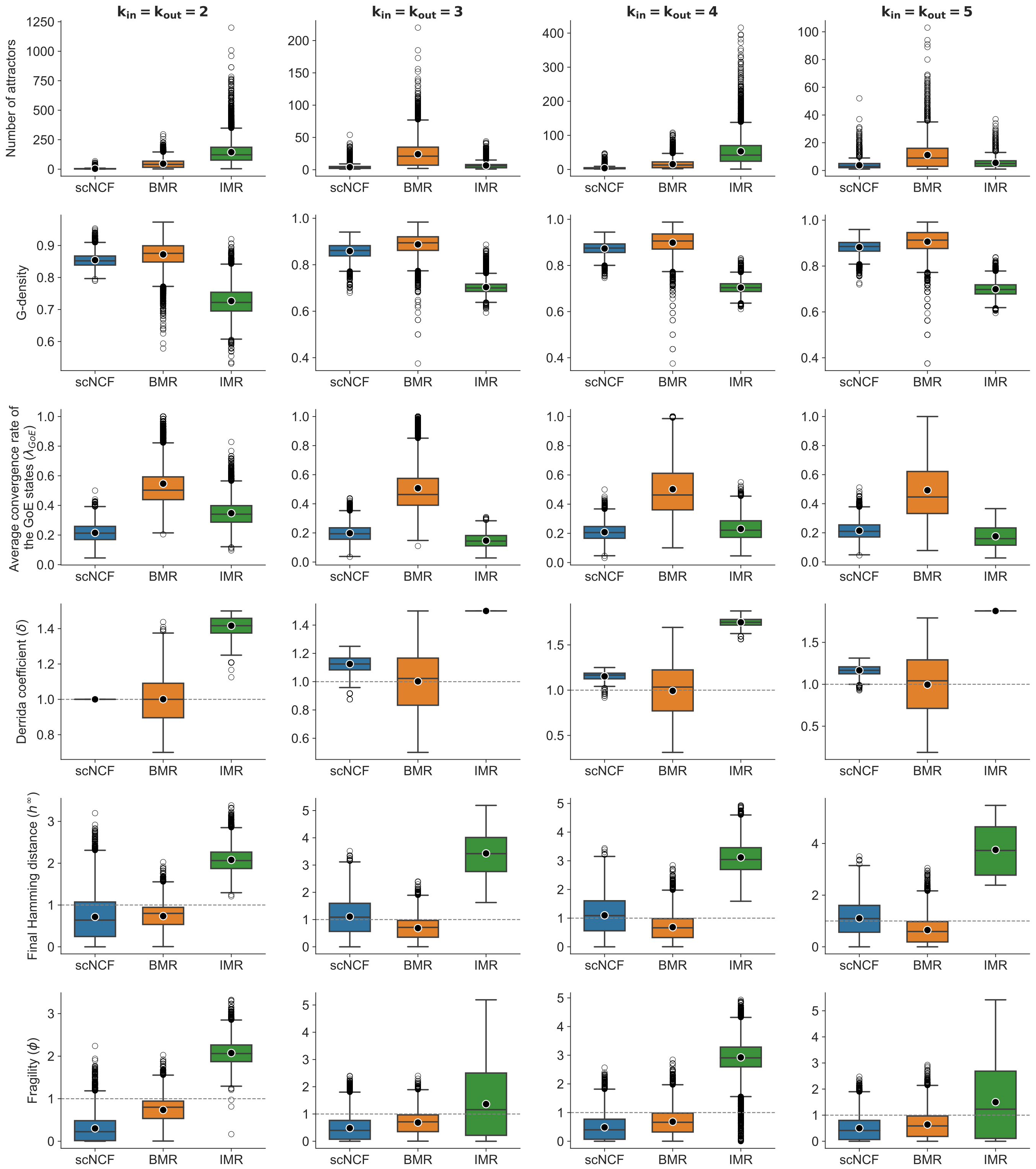


FIG. S7. **Distributions of stability measures for different subtypes of ThFs under the R-R network topology with $N = 12$.** Each row corresponds to one observable and each of the four columns represent different values of the fixed in-degree ($k_{in} = k_{out} = 2, 3, 4, 5$). In each subplot, the three boxplots present the distributions of the observables across three subtypes of ThFs: scNCFs, BMRs and IMRs. Outliers in each distribution are shown as grey circles while the black dots outlined in white denote the mean of each group.

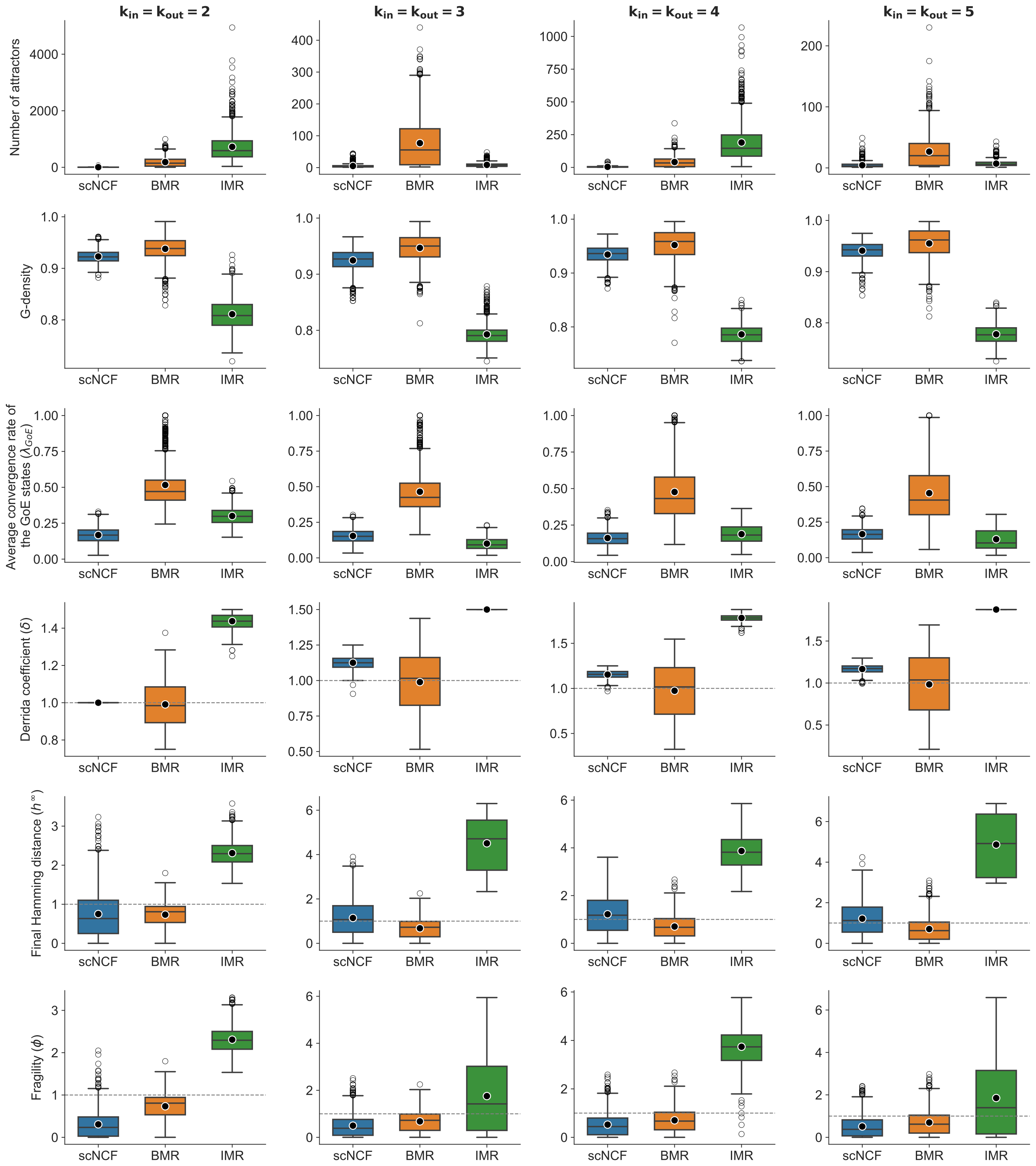


FIG. S8. **Distributions of stability measures for different subtypes of ThFs under the R-R network topology with $N = 16$.** Each row corresponds to one observable and each of the four columns represent different values of the fixed in-degree ($k_{in} = k_{out} = 2, 3, 4, 5$). In each subplot, the three boxplots present the distributions of the observables across three subtypes of ThFs: scNCFs, BMRs and IMRs. Outliers in each distribution are shown as grey circles while the black dots outlined in white denote the mean of each group.

[1] L. Raeymaekers. Dynamics of Boolean networks controlled by biologically meaningful functions. *Journal of Theoretical Biology*, 218(3):331–341, 2002.

[2] J. Aracena. Maximum Number of Fixed Points in Regulatory Boolean Networks. *Bulletin of Mathematical Biology*, 70(5):1398–1409, 2008.

[3] S. A. Kauffman. *The origins of order: self-organization and selection in evolution*. Oxford University Press, New York, 1993.

[4] M. C. Golumbic and V. Gurvich. *Read-once functions. Encyclopedia of Mathematics and its Applications*. Cambridge University Press, Cambridge, 2011.

[5] É. Lucas. Sur les congruences des nombres eulériens et des coefficients différentiels des fonctions trigonométriques suivant un module premier. *Bulletin de la Société mathématique de France*, 6:49–54, 1878.

[6] C. W. Wu. Sums of products of binomial coefficients MOD 2 and run length transforms of sequences. *Integers*, 22:A81, 2022.

[7] S. Zhang. Note on the average sensitivity of monotone Boolean functions. *preprint*, page 4, 2011.

[8] A. Biswas and P. Sarkar. Counting Unate and Monotone Boolean Functions Under Restrictions of Balancedness and Non-Degeneracy. *Journal of Integer Sequences*, 28:1–19, 2025.
